# A distinct phase of cyclin B (Cdc13) nuclear export at mitotic entry in *S. pombe*

**DOI:** 10.1101/2025.06.05.658100

**Authors:** Samir G. Chethan, Jessie M. Rogers, Drisya Vijayakumari, Wendi Williams, Vojislav Gligorovski, Sahand Jamal Rahi, Silke Hauf

## Abstract

In eukaryotes, cell division requires coordination between the nucleus and cytoplasm. Entry into cell division is driven by cyclin-dependent kinases (CDKs), which need a cyclin binding partner for their activity. In *Schizosaccharomyces pombe* (fission yeast), the B-type cyclin Cdc13 is essential and sufficient for cell cycle progression and is strongly enriched in the nucleus. Here, we show that a fraction of Cdc13 is exported from the nucleus to the cytoplasm just prior to mitosis. This export could be critical to propagate CDK activity throughout the cell. Mutating three Cdc13 nuclear localization signals (NLSs) led to precocious enrichment of Cdc13 in the cytoplasm but did not accelerate mitotic entry, indicating that the export is not sufficient to trigger entry into mitosis. The export coincides with spindle pole body integration into the nuclear envelope and may be required to coordinate nuclear and cytoplasmic signaling required for this integration. The onset and stop of Cdc13 nuclear export are remarkably abrupt, underscoring that *S. pombe* mitotic entry consists of several switch-like transitions over the course of minutes. Our findings add another instance to the various cyclin nuclear transport events known to occur at critical cell cycle transitions throughout eukaryotes.

## Background

The eukaryotic cell cycle is characterized by distinct phases (G1, S, G2, M) with rapid transitions from G1 to S, and G2 to M. The key drivers of the cell cycle are cyclin-dependent kinases (CDKs), which require binding to a cyclin protein for their activity [1–3]. In *Schizosaccharomyces pombe* (fission yeast), the cyclin-dependent kinase CDK1 (*S.p.* Cdc2) is essential for cell cycle progression and can pair with several cyclins [4]. The B-type cyclin Cdc13 is the only one of these cyclins that is essential for viability, and Cdc13 is sufficient to execute the cell cycle in the absence of the other Cdc2-binding cyclins [5,6]. Cdc13 accumulates during the S and G2 phases of the cell cycle and is degraded by APC/C-mediated proteasomal degradation late in mitosis and G1. In all cell cycle phases where Cdc13 is present, it is strongly enriched in the nucleus [7–12].

The localization of cell cycle cyclins (and their CDK binding partners) is often dynamic [13]. *S. pombe* Cdc13–in addition to its nuclear localization–binds to spindle pole bodies (SPBs) starting in late G2 phase and to the mitotic spindle during mitosis [12,14–19]. Cdc13 can also become enriched in the nucleolus at times [20–23]. In mammalian cells, cyclin A2 is mostly nuclear but becomes enriched in the cytoplasm after S phase and at centrosomes (equivalent to yeast spindle pole bodies) in early mitosis [24–31]. Mammalian cyclin B1 is mostly cytoplasmic but also localizes to centrosomes and is rapidly imported into the nucleus at entry into mitosis [24,25,32–34]. Cyclin B1’s nuclear import propagates CDK1 activity into the nucleus [34,35], and creates a spatial positive feedback loop, supporting a rapid and irreversible transition from G2 phase to mitosis [36,37]. In mitosis, cyclin B1 also localizes to spindle microtubules and kinetochores [24,32,36,38–41].

Here, we report that a fraction of *S. pombe* Cdc13 becomes exported from the nucleus just prior to mitosis, which has recently also been described by Kapadia and Nurse [42]. We find that Cdc13 export starts concomitant with Polo-like kinase (Plo1) enrichment at SPBs and stops when spindle pole bodies separate and the mitotic spindle forms. Along with other observations [42–48], our findings indicate that *S. pombe* mitotic entry is a fast sequence of at least three distinct switch-like events, likely coupled to the integration of the SPBs into the nuclear envelope.

## Results

### Cdc13 and Cdc2 export from the nucleus prior to degradation of Cdc13 in mitosis

When performing live-cell imaging of Cdc13-sfGFP, we observed a drop in nuclear concentration (**Fig. 1A**, black arrowheads) that occurred just prior to mitosis and distinctly prior to the degradation of Cdc13 in late mitosis (**Fig. 1A**, gray arrowheads). This drop is too subtle to be observed in population data when cells are arranged by size (**Fig. S1**) [18] but has also been observed by others using live-cell imaging, either with the same or another Cdc13 tag [42,48]. The drop in nuclear concentration was initially interpreted as reflecting movement of Cdc13 to the spindle pole body (SPB) [48]. While SPB localization is visible in this period (**Fig. 1B**, arrowheads), we also detect a more general increase in cytoplasmic Cdc13 concentration in the same period, whereas the cellular concentration of Cdc13 stayed comparatively constant (**Fig. 1A-C**). Highly similar measurements were made by Kapadia and Nurse [42]. When we plotted the total amounts of Cdc13 in the cell, nucleus, and cytoplasm, rather than the concentration, the cytoplasmic amount increased faster than the total amount and concomitant with a decrease in the nuclear amount (**Fig. 1D**). This indicates that a fraction of Cdc13 is exported from the nucleus at this time, prior to Cdc13 degradation in late mitosis (**Fig. 1E**). (Note that what we call “export” could either be export, or less efficient import of Cdc13 that cycles between nucleus and cytoplasm.)

**Figure 1.**
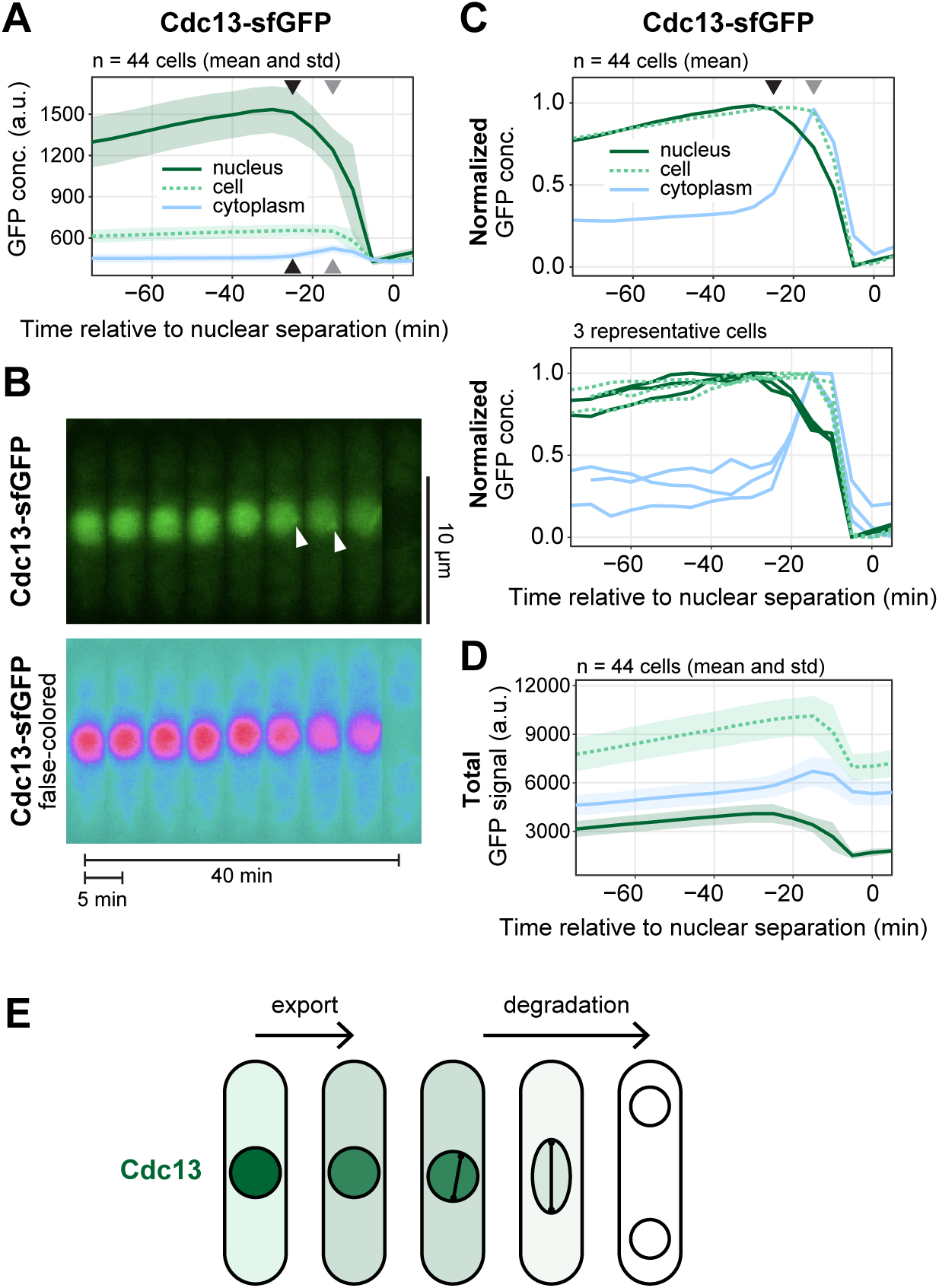
A fraction of Cdc13 exports from the nucleus prior to spindle formation. **(A)** Live-cell imaging of Cdc13-sfGFP, expressed from its endogenous genetic locus. Mean (line) and standard deviation (shaded area) of GFP concentrations in different compartments across 44 cells. Images were taken every 5 min. Curves were aligned to nuclear separation in anaphase (time 0). Cells were segmented based on the brightfield image, nuclei were segmented using an NLS-tdTomato signal. GFP concentrations are calculated as integrated signal per area. The cytoplasmic concentration was calculated from the difference in integrated signals between cell and nucleus divided by the area difference. Black arrowheads mark the approximate start of Cdc13 export from the nucleus, gray arrowheads mark the approximate start of Cdc13 degradation. Kymograph of an exemplary cell from the experiment in (A). Arrowheads point to localized Cdc13 signals, likely colocalizing with SPBs. **(C)** GFP concentrations were normalized to the minimum and maximum signal within one cell cycle for each cell to better demonstrate the changes in whole cell and cytoplasmic GFP concentration. The mean across 44 cells (top) and three exemplary cells (bottom) are shown. **(D)** The integrated signal in different regions (mean and standard deviation) is shown rather than the concentration. (E) Schematic of Cdc13 nuclear export prior to spindle pole separation and Cdc13 degradation at the metaphase-to-anaphase transition.

Cdc13 binds Cdc2 and is required for the nuclear localization of Cdc2 [7,8,12]. We found that Cdc2 nuclear concentration also started to drop prior to mitosis, accompanied by an increase in cytoplasmic concentration (**Fig. 2A**, black arrowheads; **Fig. S2**). An additional, further decrease in nuclear and increase in cytoplasmic Cdc2 concentration is observed later, when Cdc13 is degraded and no longer available as a binding partner (**Fig. 2A**, gray arrowheads; **Fig. S2**). This suggests that Cdc13 and Cdc2 are exported from the nucleus as a complex, and, therefore, that CDK1 activity may spread to the cytoplasm during this time. As a control, another nuclear-enriched protein (Mad3, a spindle assembly checkpoint protein) did not show a drop in nuclear concentration during that same period (**Fig. 2A, S2**), indicating that the export is actively regulated rather than representing unspecific leakiness of the nuclear envelope. Leakiness may have occurred because the *S. pombe* SPB integrates into the nuclear envelope during this time, which requires nuclear envelope fenestration [49,50].

**Figure 2.**
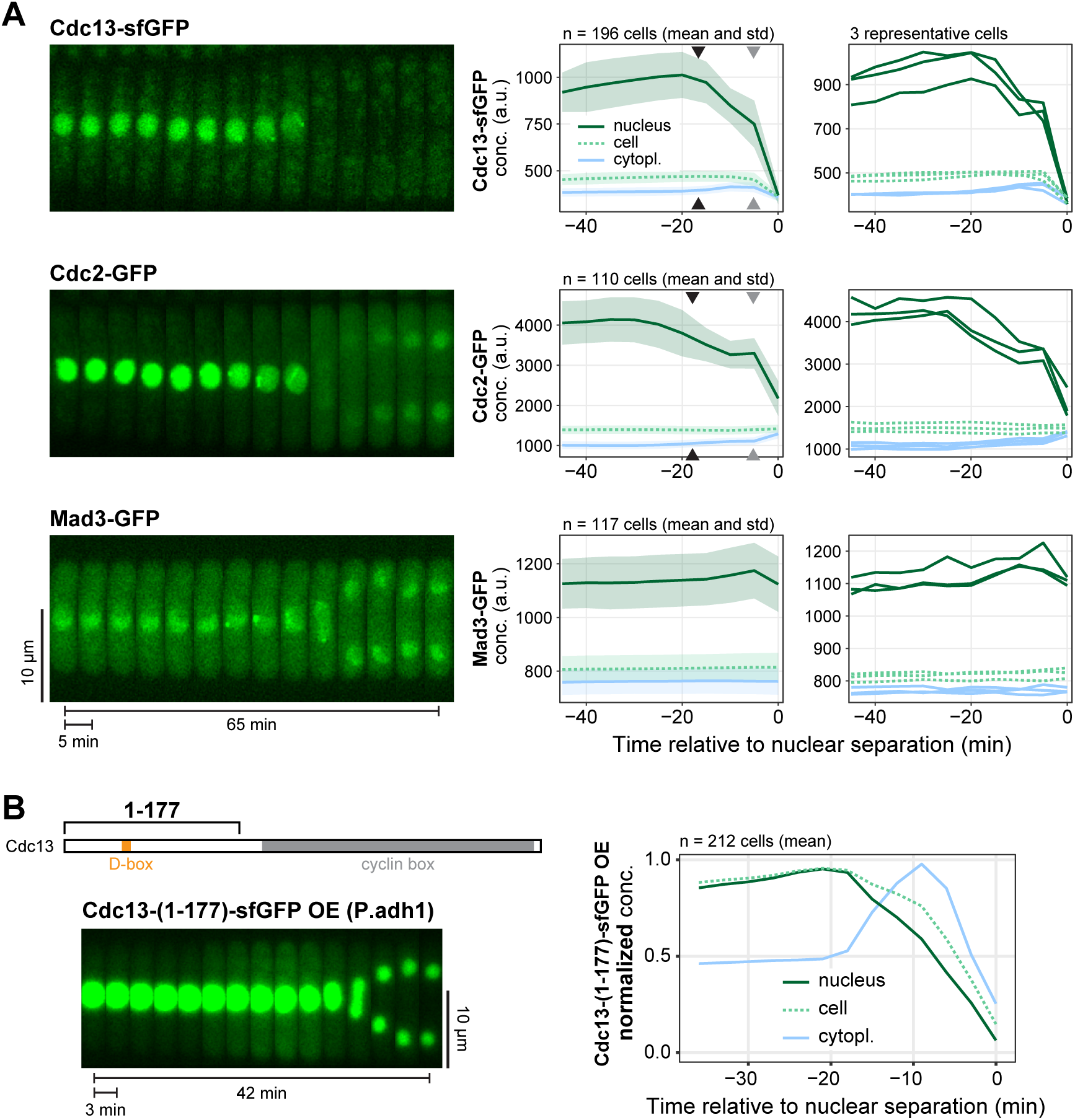
Cdc2 exports from the nucleus along with Cdc13, but Cdc13 can export without Cdc2. **(A)** Live-cell imaging of Cdc13-sfGFP, Cdc2-GFP, and Mad3-GFP, similar to Fig. 1, all expressed from their endogenous genetic loci. Images were taken every 5 min. Kymographs from exemplary cells (left), mean and standard deviation of the GFP concentrations (middle), and GFP concentrations from three exemplary cells each (right). Black arrowheads mark the approximate start of Cdc13 and Cdc2 export from the nucleus, gray arrowheads mark the approximate start of Cdc13 degradation. Normalized GFP concentrations are shown in Fig. S2. **(B)** Live-cell imaging of an N-terminal Cdc13 fragment (amino acids 1–177), C-terminally tagged with sfGFP and overexpressed (OE) from the strong *adh1* promoter at the *leu1* locus. Images were taken every 3 min. Imaging and analysis otherwise as in (A). Kymograph (left), and GFP concentration normalized to the minimum and maximum signal within one cell cycle (right).

Even though Cdc13 and Cdc2 may export as a complex, the export of Cdc13 does not require Cdc2, since the N-terminal unstructured region of Cdc13, Cdc13-(1-177), which is not expected to interact with Cdc2 [38,51,52], shows the same brief period of nuclear export (**Fig. 2B**). (Note that Cdc13-(1-177) is overexpressed in this experiment, making the increase in cytoplasmic concentration more obvious).

### Cdc13 export takes place prior to SPB separation and is concomitant with Plo1 enrichment at SPBs

To better assess the timing of Cdc13 export, we co-labeled the centromere of chromosome 1 with tdTomato, took images every 15 seconds, and aligned the measurements to sister chromatid separation in anaphase (**Fig. 3, Fig. S3**). Nuclear export of Cdc13 began about 15 min prior to anaphase and, remarkably, ceased about 7 min prior to anaphase, which corresponds to the time when spindle poles separate. Another APC/C substrate, securin (*S.p.* Cut2), showed a different pattern: while Cdc13 was exporting, Cut2 nuclear concentration stayed constant, but it imported into the nucleus starting at SPB separation (**Fig. 3**). This confirms that Cdc13 nuclear export is not a consequence of nuclear leakiness and indicates two different phases of nuclear transport regulation at entry into mitosis: before and after spindle pole separation.

**Figure 3.**
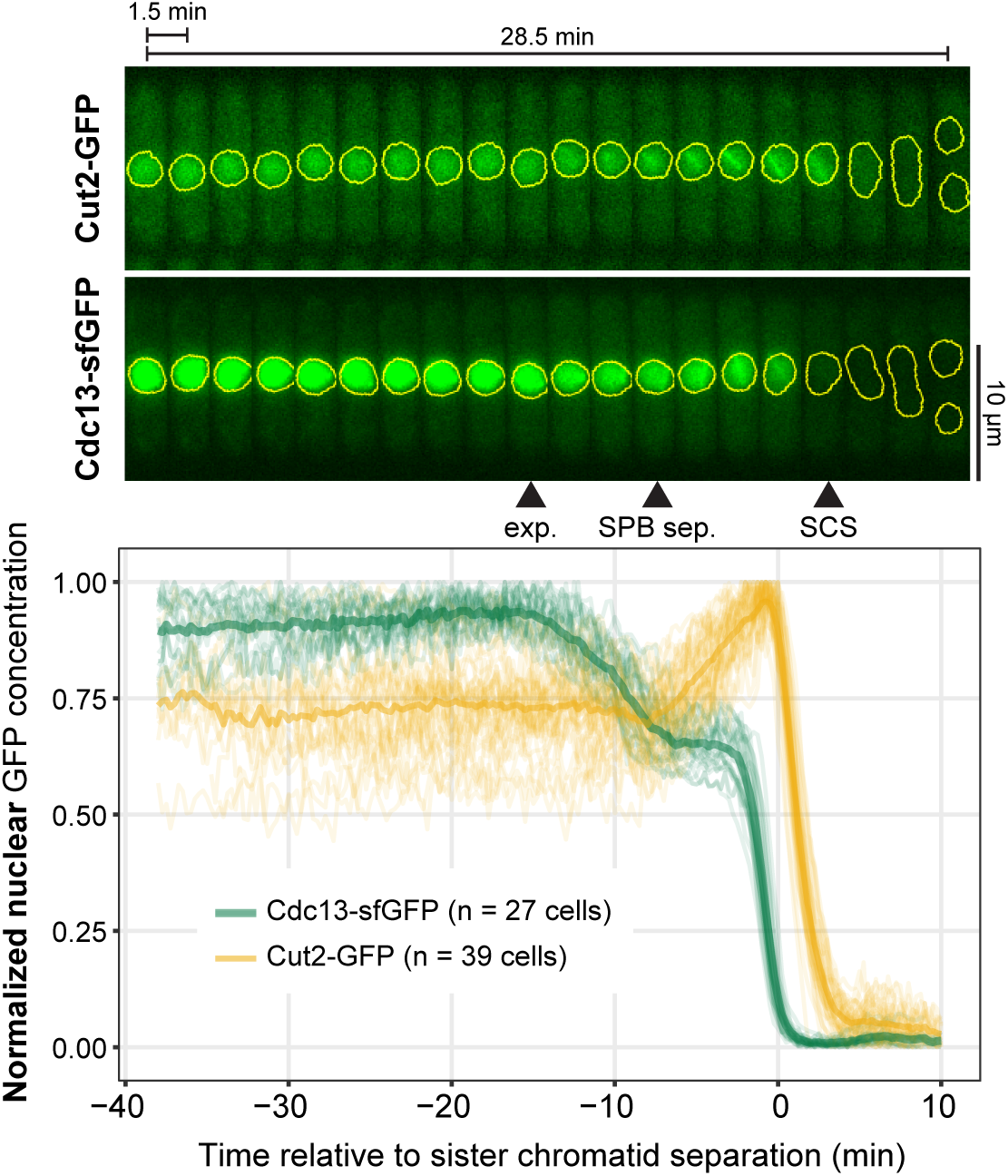
Cdc13 stops exporting from the nucleus at spindle formation. Live-cell imaging of Cdc13-sfGFP and Cut2-GFP, both expressed from their endogenous genetic locus. Images were taken every 15 sec; the kymographs from exemplary cells (top) only show every 6th image, yellow shapes indicate the area segmented as nucleus. Arrowheads mark the approximate positions of start of Cdc13 nuclear export (exp.), SPB separation (SPB sep.), and sister chromatid separation (SCS). The centromeres of chromosome 1 were labelled with tdTomato (Fig. S3), and nuclear GFP concentration curves were aligned to sister chromatid separation in anaphase (time 0). Nuclear concentrations (bottom) of individual cells (n = 27 for Cdc13, n = 39 for Cut2) as thin lines, their mean as thick line. Concentrations were normalized to the maximum and minimum for each cell. To make the strains as similar as possible, Cut2 is tagged with non-fluorescent GFP in the Cdc13-sfGFP strain, and Cdc13 is tagged with non-fluorescent sfGFP in the Cut2-GFP strain. Exemplary single cells are shown in Fig. S4.

At this higher temporal resolution, it became apparent that the onset and stop of nuclear export are rapid events, observable by the abrupt slope change both in the averaged data (**Fig. 3**) as well as in single cells (**Fig. S4**). The transition to decreasing nuclear Cdc13 concentration happens within about a minute and the transition from export to its cessation is typically even more abrupt (**Fig. S4**). This suggests that the onset and stop of nuclear export are switch-like transitions and require a type of molecular regulation that can implement such rapid change.

To position the onset of Cdc13 nuclear export relative to other events, we looked at Plo1 localization. Plo1 (Polo or Plk1 in other organisms) is a key kinase in the regulation of mitotic entry in both mammalian cells and *S. pombe* [53–57]. In *S. pombe*, Plo1 becomes enriched at SPBs in late G2, just prior to mitosis, and SPBs serve as a signaling hub for mitotic entry regulation involving Plo1 and Cdc2 [12,17,19,57–61]. We imaged Plo1-mCherry together with Cdc13-sfGFP. In these experiments, Cdc13-sfGFP was expressed from the endogenous locus either under the endogenous *cdc13* promoter or overexpressed under the *adh1* promoter (**Fig. 4A**). These experiments confirmed that Cdc13 nuclear export takes place prior to SPB separation and showed that its start is concomitant with the enrichment of Plo1 at SPBs and stops with SPB separation (**Fig. 4A,B**). Fission yeast SPBs are located outside the nuclear envelope during interphase and need to integrate into the nuclear envelope for mitotic spindle formation [49,50,62,63]. The coincidence in timing suggests that Cdc13 export is important for SPB integration.

**Figure 4.**
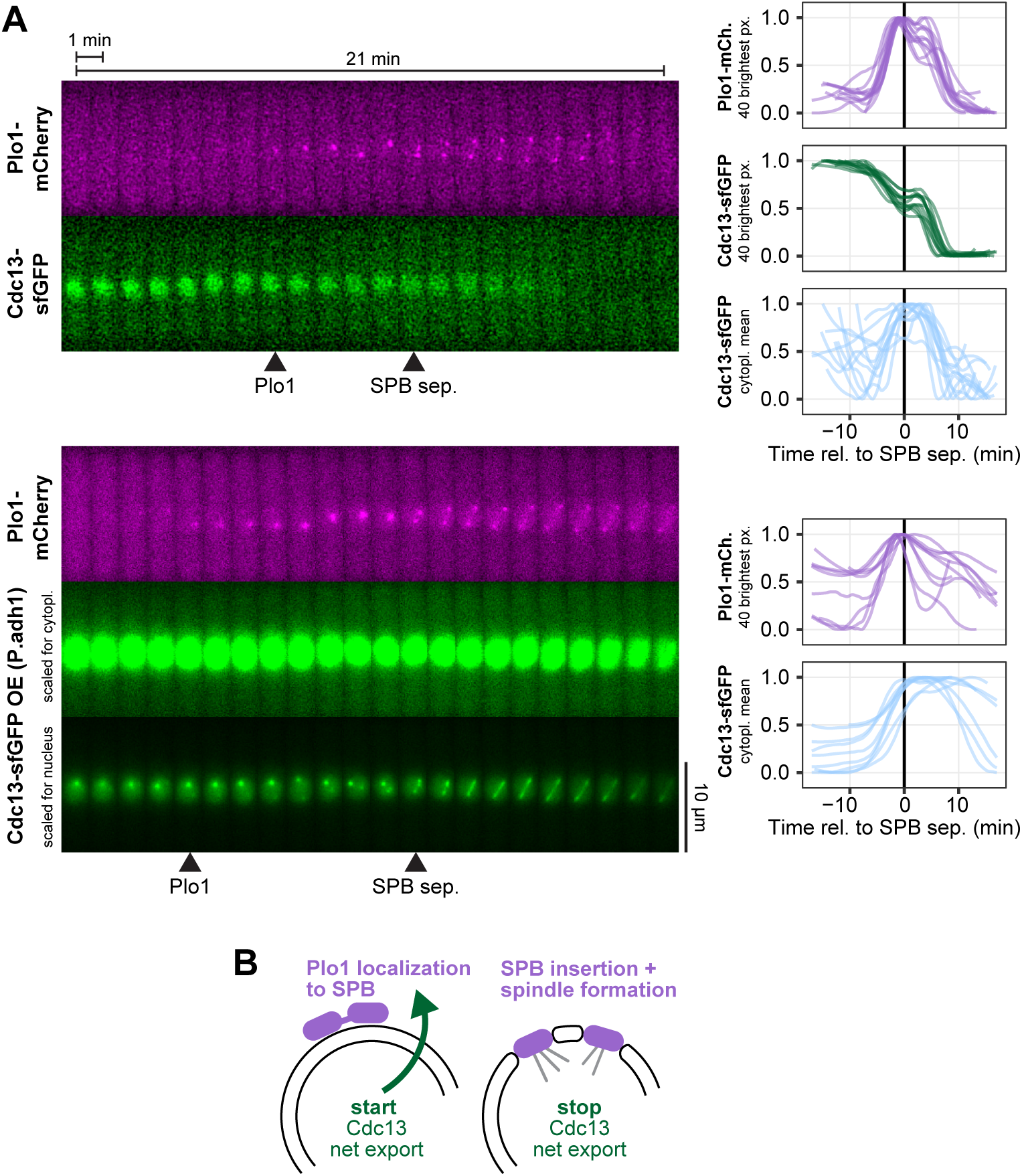
Cdc13 nuclear export is concomitant with Plo1 enrichment at the spindle pole bodies. **(A)** Live-cell imaging of Plo1-mCherry and Cdc13-sfGFP, both expressed from their endogenous genetic locus. Cdc13-sfGFP expressed from the *cdc13* promoter (top) or the strong *adh1* promoter (bottom). Images were taken every 15 sec; the kymographs from exemplary cells (left) only show every 4th image. Graphs on the right show concentrations from individual cells normalized to the minimum and maximum signal recorded for each cell (n = 12 and n = 9 cells, respectively). Curves were aligned to SPB separation (time 0). Since the strains did not express a nuclear marker, the concentration of Cdc13 in the nucleus was estimated from the mean of the 40 brightest pixels in the cell. This was not possible for the overexpression strain due to strong accumulation of the overexpressed Cdc13 at a dot-like spot within the nucleus, likely the rDNA (bottom panel in bottom kymograph). This signal was so strong that it also bled through into the mCherry channel (seen as very weak dot-like signal in the images just before Plo1-mCherry starts to concentrate at the spindle pole bodies). The mean of the 40 brightest pixels was also used to quantify the accumulation of Plo1-mCherry at the spindle pole bodies. GFP concentration in the cytoplasm (cytopl. mean) was measured from two manually placed regions on either side of the nucleus. Arrowheads on the kymographs mark the approximate positions of the start of Plo1 enrichment at the SPB region (Plo1) and SPB separation (SPB sep.). Cdc13 overexpression (∼ 20x, bottom panel) can cause delays in mitosis, leading to the persistence of Cdc13-sfGFP beyond 10 min seen for some cells quantified in the bottom panel. **(B)** Schematic illustrating that nuclear export of Cdc13 starts concomitantly with the enrichment of Plo1 at SPBs and stops concomitantly with SPB separation.

### Movement of Cdc13 from nucleus to cytoplasm is not sufficient for mitotic entry

To address whether the export of Cdc13 from the nucleus influences the timing of entry into mitosis, we sought to precociously enrich Cdc13 in the cytoplasm. The N-terminal unstructured region of Cdc13 contains three candidate nuclear localization signals (NLSs) (**Fig. 5A**, **Fig. S5**) [8]. We mutated these and expressed the mutant constructs under the endogenous regulatory sequences from an exogenous locus, leaving the endogenous *cdc13* intact. While mutation of individual NLSs did not prominently affect Cdc13 nucleocytoplasmic distribution (**Fig. 5B,C**, and not shown), combining mutations in all three NLSs led to a reduction in nuclear enrichment (**Fig. 5B,C**, NLS1-2-3 mutant). We initially left two lysine residues that are part of the predicted bi-partite NLS1 intact because they are directly adjacent to the destruction box (D-box), and we assumed they may be important targets for ubiquitination and therefore essential for the rapid proteasome-mediated degradation of Cdc13 in late mitosis. However, mutation of these residues still allowed for degradation of Cdc13 at the end of mitosis while further impairing nuclear enrichment (**Fig. 5B,C, 6A**, **S6A,B**, NLS-KK mutant). In live-cell imaging, a prominent cytoplasmic pool of Cdc13-NLS-KK was observed throughout the cell cycle (**Fig. 6A, S6B**). Despite the precocious and strong enrichment of Cdc13-NLS-KK in the cytoplasm, the length of the cell cycle was not shortened in this strain (**Fig. 6B**), indicating that Cdc13 nuclear export is not sufficient to trigger mitotic entry. This situation is similar to that of mammalian cyclin B1, which rapidly imports into the nucleus at entry into mitosis, but in cell lines, a constitutive or precocious nuclear localization of cyclin B1 does not accelerate entry into mitosis [64–67].

**Figure 5.**
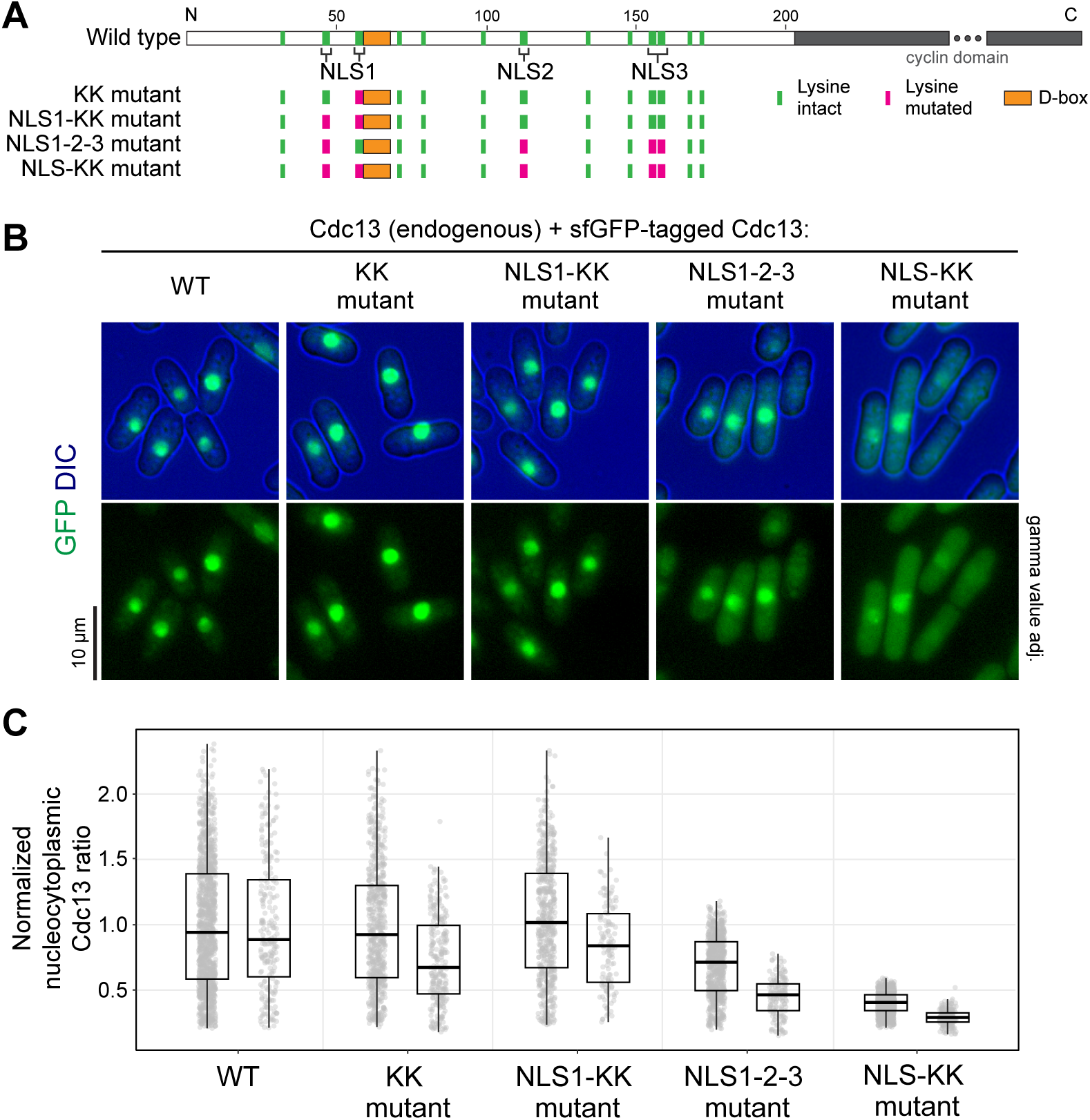
Mutation of three Cdc13 nuclear localization signals causes accumulation of Cdc13 in the cytoplasm. **(A)** Schematic depicting the location of lysine residues and the D-box within the Cdc13 protein disordered region. Residues mutated in respective constructs are highlighted in magenta. **(B)** Representative images of asynchronous cultures of cells expressing wild-type or mutant versions of Cdc13-sfGFP expressed from the exogenous *leu1* locus under the endogenous *cdc13* promoter. In images showing GFP only, the gamma values were adjusted to better illustrate the range of signals. **(C)** Quantification of the nucleocytoplasmic ratio of Cdc13 concentration for wild-type Cdc13 and NLS mutant versions. Two independent experiments, imaged on different microscopes. Nucleocytoplasmic Cdc13 ratios for each cell were normalized to the mean nucleocytoplasmic ratio of Cdc13 in wild-type cells within each respective experiment. n > 550 and >120 cells per strain in experiment 1 and 2, respectively. Individual cells (circles) and summary data; box: median and quartiles; whiskers: 1.5-times interquartile range.

**Figure 6.**
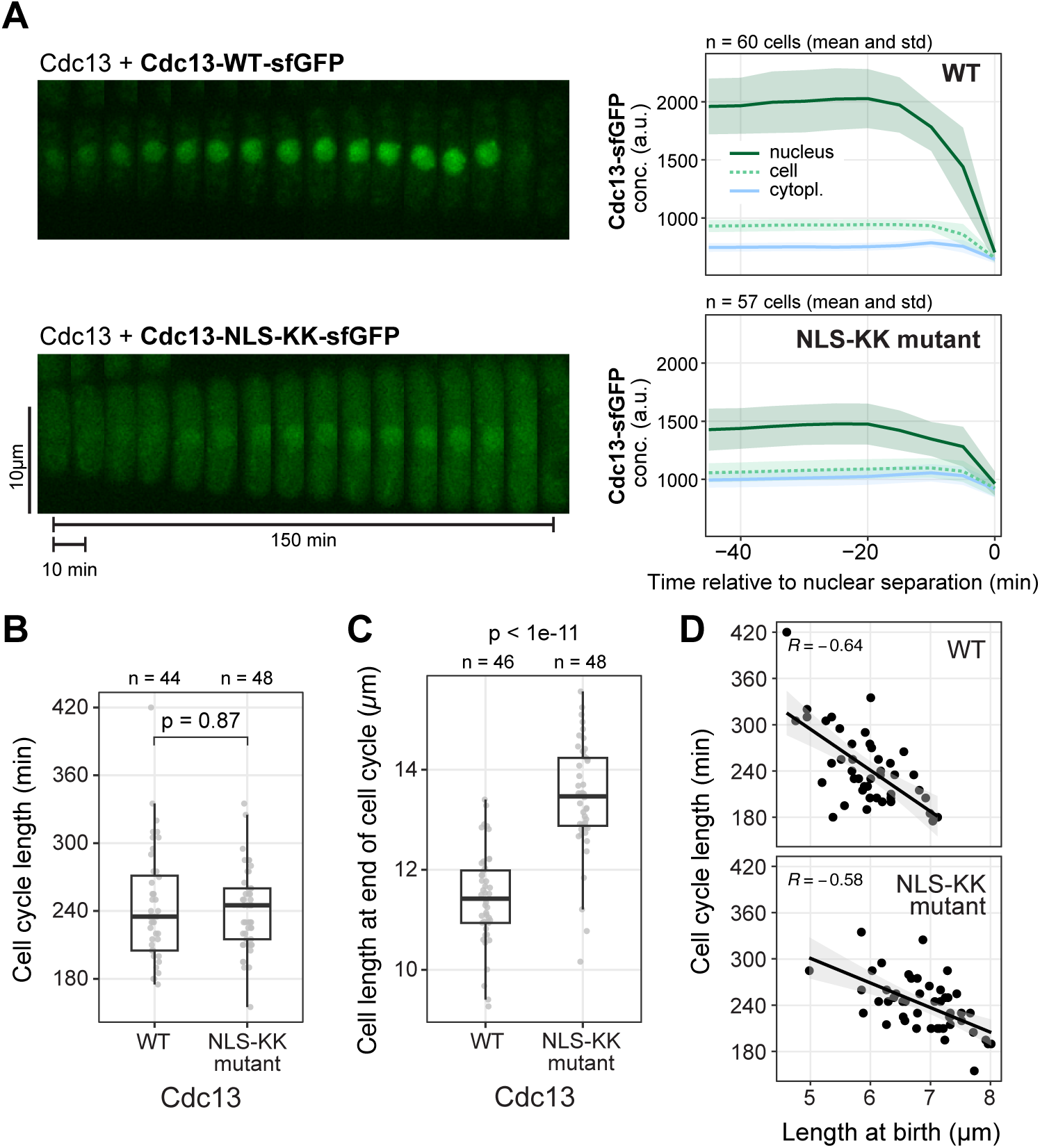
Cdc13 nuclear export is not the rate-limiting step for entry into mitosis. **(A)** Live-cell imaging of wild-type (WT) or NLS-KK mutant Cdc13, similar to Fig. 1, except that the fluorescently tagged versions were expressed from the exogenous *leu1* locus under the *cdc13* promoter and untagged, wild-type *cdc13* is expressed from its endogenous locus. Images were taken every 5 min. Kymographs from exemplary cells (left), mean and standard deviation of the GFP concentrations (right). Plots for the normalized and total GFP signal are available in Fig. S6. **(B,C)** Time from one cell division to the next (B) and cell length in the time frame prior to cell division (C) from the live-cell imaging experiment in (A). Individual cells (circles) and summary data; box: median and quartiles; whiskers: 1.5-times interquartile range. p-values from Wilcoxon rank sum test with continuity correction. **(D)** Cell length at birth versus length of the subsequent cell cycle from the live-cell imaging experiment in (A). Individual cells (circles) and linear regression. R, Pearson correlation coefficient.

Consistent with the unaltered cell cycle length, we found that strong enrichment of Plo1 at SPBs remained restricted to ∼ 8 min prior to SPB separation in cells expressing Cdc13-NLS-KK (**Fig. 7, S7**). Plo1 enrichment still coincided with Cdc13 nuclear export. Nuclear export was detectable as a drop in nuclear signal in the Cdc13-NLS-KK mutant despite its lowered nucleocytoplasmic ratio (**Fig. 7**, **S7**). This suggests that the change in nucleocytoplasmic Cdc13 distribution prior to mitosis is not a consequence of blocking NLS function—at least not of those NLSs that are mutated in Cdc13-NLS-KK. What mediates the remaining nuclear enrichment of Cdc13-NLS-KK is unknown. Cdc13 does not contain other classical NLSs. Nuclear accumulation of Cdc13 therefore may partially rely on non-canonical mechanisms such as direct binding to importin-β, as is the case for other cyclins [27,68–70].

**Figure 7.**
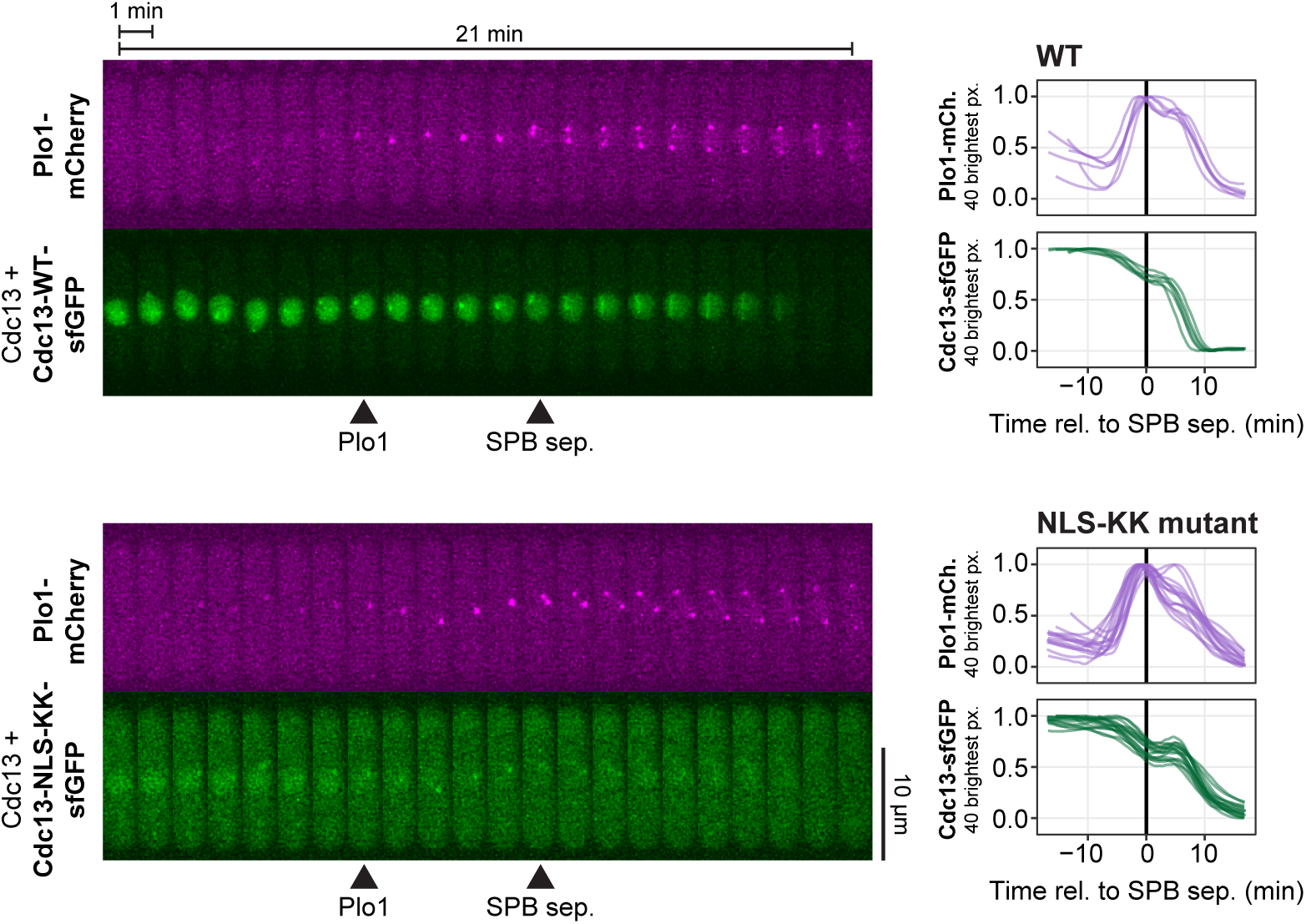
Plo1 enrichment at spindle pole bodies is not accelerated by cytoplasmic Cdc13 in the NLS-KK mutant. Live-cell imaging of Plo1-mCherry, expressed from its endogenous locus, and sfGFP-tagged wild-type Cdc13 or NLS-KK mutant, expressed from the exogenous *leu1* locus under the *cdc13* promoter. Untagged, wild-type *cdc13* is expressed from its endogenous locus. Images were taken every 15 sec; the kymographs from exemplary cells (left) show every 4th image. Arrowheads on the kymographs mark the approximate positions of the start of Plo1 enrichment at the SPB region (Plo1) and SPB separation (SPB sep.). Graphs on the right show concentrations from individual cells normalized to the minimum and maximum signal recorded for the cell (n = 6 and n = 16 cells from two independent strains, respectively). Curves were aligned to SPB separation (time 0). The mean of the 40 brightest pixels in the cell was used to quantify the accumulation of Plo1-mCherry at the spindle pole bodies and obtain an estimate for the nuclear Cdc13 concentration. Plots for the non-normalized concentrations and additional kymographs are shown in Fig. S7.

Taken together, the unaltered cell cycle length and similar kinetics of Plo1 accumulation at SPBs in Cdc13-NLS-KK-expressing cells suggest that increasing CDK1 activity in the cytoplasm is insufficient to trigger mitotic entry.

While the cell cycle length of Cdc13-NLS-KK mutant-expressing cells was unchanged, the size of these cells increased (**Fig. 6C**, **S4C**). Size homeostasis (longer cell cycles in cells that are born shorter [71]) was maintained (**Fig. 6D**, **S4D**), indicating that the “set point” for size had changed without altering the regulation of size control. We attribute this to the altered distribution of Cdc13 (and therefore likely Cdc2) between the nucleus and the cytoplasm. With less nuclear CDK1 activity, cells may reach the threshold for mitotic entry only at a larger size. The alteration of cell size is consistent with observations that perturbing the nucleocytoplasmic ratio of the Cdc2 regulators Wee1 or Cdc25 alters cell size [45,72,73].

## Discussion

The nucleus of eukaryotic cells provides additional opportunities for the regulation of cellular activities but also brings about the need to coordinate nuclear and cytoplasmic events. Changes in the nucleocytoplasmic distribution of cyclins or other CDK regulators are a major theme in cell cycle regulation. Here, we show that a pool of *S. pombe* Cdc13 translocates from the nucleus to the cytoplasm at about the time when Plo1 starts to become strongly enriched at SPBs. We suggest that this is part of a sequence of events leading to the insertion of the SPBs into the nuclear envelope and, ultimately, mitotic spindle formation (**Fig. 8**). The export could serve to spread CDK1 activity into the cytoplasm, as also proposed by Kapadia and Nurse [42], who have shown, using biosensors for CDK1 activity, that CDK1 becomes active in the nucleus prior to Cdc13 export and prior to detectable CDK1 activity in the cytoplasm. The Cdc13 translocation seems functionally analogous to the movement of vertebrate cyclin B1 at entry into mitosis [34], except that the directionality is reversed (vertebrate cyclin B1 moving from the cytoplasm to the nucleus, and *S. pombe* Cdc13 moving in the other direction).

**Figure 8.**
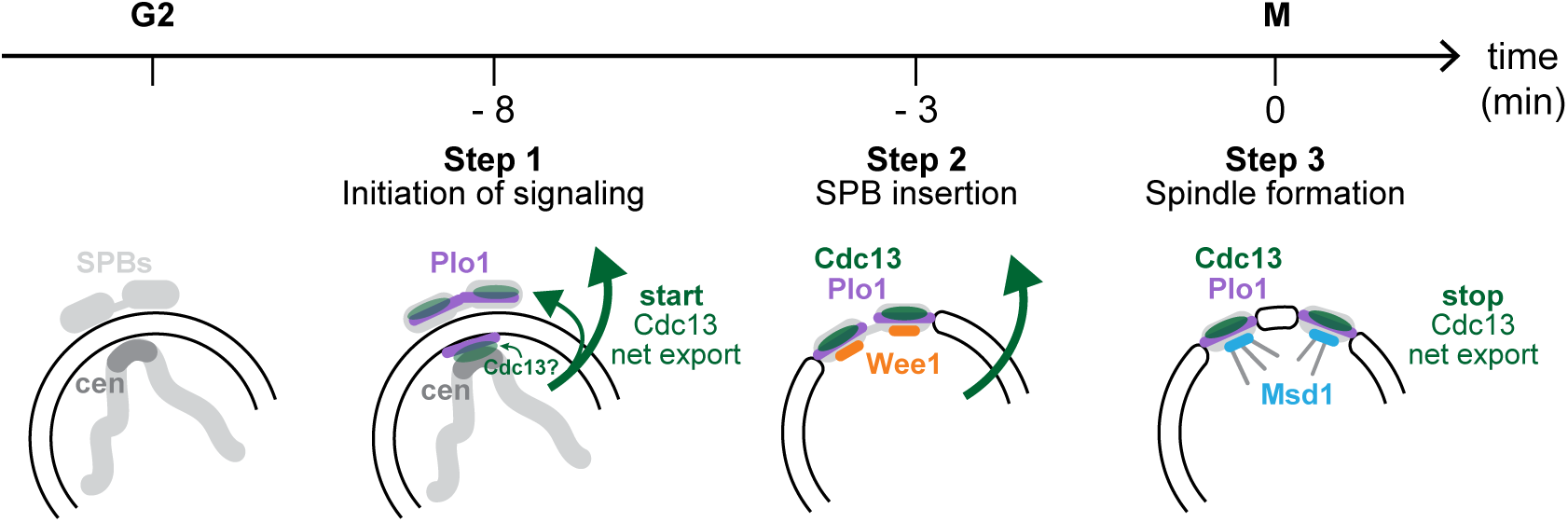
Cdc13 export is part of a stepwise entry into mitosis, coincident with SPB insertion into the nuclear envelope. Model for multiple steps at the S. pombe G2/M transition. The start of Cdc13 nuclear export coincides with Plo1 and Cdc13 enrichment at the SPB region. SPB integration into the nuclear envelope requires centromere (cen) contact with the SPB region (Fernandez-Alvarez et al., 2016; Bestul et al., 2021). Plo1 localizes both to cytoplasmic SPBs and to the nucleoplasmic side of the nuclear envelope adjacent to the SPBs (Bestul et al., 2021). The precise Cdc13 localization in this time window has not been determined. Cdc13 nuclear export could be required to bring active Cdc13/Cdc2 into contact with Plo1 at the cytoplasmic SPBs to jointly orchestrate SPB insertion into the nuclear envelope. Wee1 localizes to the SPBs distinctly later than Plo1 (Masuda et al., 2011), possibly when SPBs have become integrated into the nuclear envelope and thus become accessible to Wee1, which is enriched in the nucleus. Cdc13 export stops around the time of spindle formation. Wee1 dissociates from SPBs at this time and other factors, such as Msd1, bind. Each transition step is fast, indicating that they involve ultrasensitivity in the underlying regulation.

### The switch-like onset and stop of Cdc13 nuclear export could be implemented by phosphoregulation

The mechanism by which the change in nucleocytoplasmic distribution of Cdc13 occurs is still unknown. One possibility is post-translational regulation, such as phosphorylation, which would be dynamic enough to change nucleocytoplasmic distribution within minutes. The onset and stop of Cdc13 export are strikingly switch-like (**Fig. 3, S4**), which suggests ultrasensitivity in the underlying regulation and could, for example, be implemented by multi-site phosphorylation or enzyme saturation [74,75]. Multiple phosphorylation sites have been identified in the N-terminal, unstructured region of Cdc13 that also contains the NLSs [22,76–81] (**Fig. S5A**), but the role of most of these sites has not yet been tested. Mutation of three phosphorylated Cdc13 residues downstream of the NLSs (S177, S180, S183) did not interfere with Cdc13 localization or function [22]. It is also possible that import or export regulators are post-translationally modified. For vertebrate cyclin B1’s rapid import into the nucleus at entry into mitosis, phosphorylation of both cyclin B1 and transport effectors has been implicated [33,34,64,82–84]. In *S. pombe*, the phosphorylation of several nucleoporins increases around M-phase, which could alter nuclear transport [79,85,86]. Furthermore, reducing the dosage of several nucleoporins has been shown to increase cell size in *S. pombe*, similar to *cdc13, cdc2*, and *cdc25* mutants [87]. Whether this is related to altered Cdc13 nuclear export remains to be tested.

### Cdc13 export may propagate CDK1 activity to the cytoplasm to coordinate step-wise SPB integration

SPBs and their vertebrate counterparts, centrosomes, are considered hubs of mitotic entry regulation [53,56,88]. In *S. pombe*, the SPBs are located adjacent to, but outside, the nuclear envelope during interphase and only become integrated into the nuclear envelope at the onset of mitosis [50,62,89]. Their proper integration is required for the formation of the mitotic spindle [43,63,90,91]. Plo1, localized to SPBs, is needed for the integration of the SPBs into the nuclear envelope [47,60,92,93], and Cdc2, in turn, is required for Plo1 localization and activity at SPBs [59,61,94,95]. We consider it plausible that Cdc2/Cdc13 export from the nucleus to the cytoplasm is required to initiate SPB integration into the nuclear envelope by co-positioning Plo1 and Cdc13/Cdc2 at the cytoplasmic SPBs (**Fig. 8**). Contact between centromeres and the nuclear envelope beneath the SPBs is also required for SPB insertion, and this signaling has also been suggested to involve Plo1 and possibly Cdc2 [46,47]. Hence, both nuclear and cytoplasmic Cdc2 activity could be important, and Cdc13 nuclear export may be required to provide them both. Interestingly, mutation of the hydrophobic patch of Cdc13 has been shown to prevent not only Cdc13 enrichment in the SPB region [17,19], but also Cdc13 nuclear export [42], which suggests positive feedback between localized CDK1 activity at the SPB region and Cdc13 export. The binding partner for the hydrophobic patch has not been identified, and it is unclear whether this reflects a defect in a nuclear or cytoplasmic interaction.

The start of Cdc13 export coincides with Plo1 enrichment at the SPB region and its stop with SPB separation (**Fig. 4**). Live-cell imaging of mitotic entry regulators by Masuda and colleagues [45] has uncovered one additional switch-like step between the enrichment of Plo1 at SPBs and SPB separation: the CDK1-inhibiting kinase Wee1 and the kinesin Cut7 become enriched at SPBs about 3 – 4 min prior to SPB separation [45], clearly after the start of Plo1 enrichment at about 6 – 8 min prior to SPB separation (**Fig. 8**). We suggest that the first wave of binding (Plo1 and Cdc13) corresponds to preparing the SPB for insertion, and the second wave (Wee1 and Cut7) corresponds to insertion into the nuclear envelope, which makes the SPB accessible to nuclear proteins such as Wee1. At SPB separation, Wee1 is removed from SPBs, and the microtubule-anchoring protein Msd1 binds [45]. This final step coincides with the cessation of Cdc13 export. Thus, *S. pombe* mitotic entry consists of at least three switch-like transitions, tightly linked to SPB events and bracketed by Cdc13 nuclear export. We propose that the export is required to provide both nuclear and cytoplasmic activities for SPB integration.

## Materials and Methods

### Strains and growth

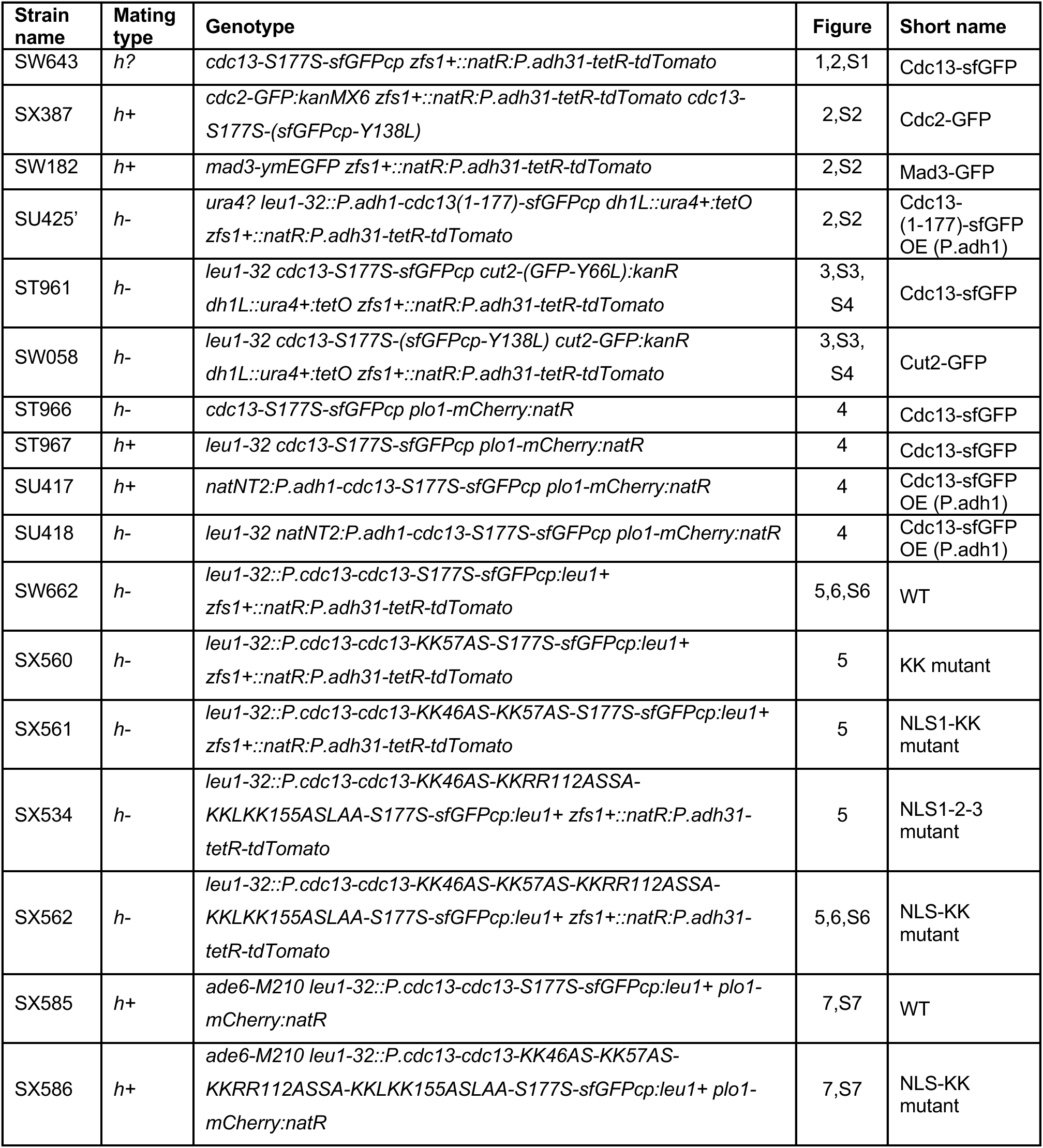

Strains with *zfs1+::natR:P.adh31-tetR-tdTomato*, *dh1L::ura4+:tetO, cdc13-S177S-sfGFPcp, P.adh1-cdc13-S177S-sfGFPcp*, *cdc2-GFP*, *mad3-ymEGFP*, *cut2-GFP*, and *plo1-mCherry* have been described previously [96–99]. Strains expressing—from an exogenous locus—only the N-terminus of Cdc13 (*cdc13(1-177)-sfGFPcp*) or Cdc13 NLS mutants were generated by cloning the respective fragments or mutant versions into a pDUAL vector [100] and integrating the vector at the *leu1* locus. The proper sequence of the vectors was confirmed by Sanger or Nanopore whole-plasmid sequencing. Some strains have a Y66L mutation introduced into GFP, which makes GFP non-fluorescent. This allows us to visualize other proteins with GFP while maintaining the tag for comparability with other strains. In circularly permuted sfGFPcp, Y138 (LT<u>Y</u>GV) is the residue corresponding to Y66 (FT<u>Y</u>GV) in canonical GFP.

### Growth conditions

Cells were grown in Edinburgh minimal medium (EMM, MP Biomedicals, 411003) at 30 °C to a concentration of 8 x 10^6^ – 1.5 x 10^7^ cells/mL. Leucine (0.2 mg/mL) or adenine (0.15 mg/mL) was added when required. Preconditioned medium (made by filtering EMM cultures) was added when cultures were diluted to low densities.

### Time-lapse imaging

Time-lapse imaging was conducted on a DeltaVision Elite microscope equipped with an Olympus 60x/1.42 Plan-APO oil objective, LED illumination, and a PCO edge sCMOS camera. Cells were kept at 30 °C for the duration of imaging (EMBL environmental chamber). Cells were either mounted in µ-Slide 8 well glass bottom chambers (Ibidi, 80827) coated with lectin (50 µg/mL; Sigma-Aldrich, L1395) or loaded into microfluidics chambers (Millipore-Sigma, Y04C or Y04T) that had been washed with EMM and prewarmed to 30 °C. Media flow for the microfluidics chambers was controlled using a CellAsic ONIX2 microfluidics system. EMM supplemented with 50 % preconditioned medium was perfused at 2 psi. After being loaded into the viewing chamber of the microfluidics plate, cells were left to acclimate for 3 hours before starting imaging. When imaging in µ-Slide chambers, cells were kept on the 30 °C microscope stage for 15 min before starting imaging. Brightfield images were taken at the bottom or central slice of each field of view and fluorescence images were acquired using “optical axis integration” (sum projection) over a 3.6 μm Z-distance. Fluorescence images in experiments shown in Figs. 2, 4, 6, and 7 were additionally deconvolved using SoftWoRx (GE Healthcare) software with three cycles of the ratio method (conservative).

### Imaging of asynchronous cultures

Asynchronous cultures were either imaged at 30 °C on a DeltaVision Elite microscope (see above), or at room temperature (∼ 22 °C) on a Zeiss AxioImager M1 equipped with Xcite Fire LED illumination (Excelitas), a Zeiss Plan-APO 63×/1.4 oil objective, and an ORCA-Flash4.0LT sCMOS camera (Hamamatsu). Cells were pelleted at 3,300 rcf for 1 minute, mounted from the pellet onto slides, covered with #1.5 glass coverslips, and immediately imaged. Z-stacks were collected and the slice corresponding to the midplane of the cell was used for the quantification of nucleocytoplasmic ratios.

### Image analysis

For the experiments in Figures 1, 2, 5, 6, S2, and S6, brightfield images were used to segment individual cells using the YeaZ neural network with custom weights [101]. Masks were manually edited using the YeaZ graphical user interface as needed. Cells partially out of frame were eliminated from the analysis. A custom Fiji [102] script was used to load corrected YeaZ cell masks as regions of interest (ROIs). Nuclei were segmented in Fiji using tdTomato-NLS fluorescence and Otsu thresholding. A Gaussian blur was applied to smooth the nuclear edges. After manual checking and correction, nuclear ROIs were assigned to cellular ROIs based on position; and size and fluorescence intensity were quantified. Cytoplasmic area and integrated intensity were calculated by subtracting nuclear area and integrated intensity from the respective whole cell measurements. To determine nucleocytoplasmic ratios, the fluorescence intensity outside cells, measured as the mean from several manually drawn ROIs, was subtracted as background. (Note that this does not take autofluorescence into account, which is difficult to quantify. Both nuclear and cytoplasmic concentrations are therefore slightly overestimated.) To calculate nucleocytoplasmic ratios, nuclear GFP concentration (integrated signal by area) was divided by the respective cytoplasmic concentration and normalized within each experiment to the mean nucleocytoplasmic ratio of cells expressing wild-type Cdc13-sfGFP.

The experiment in Figure 4 lacked a brightfield image for cell segmentation and a fluorescent marker for nuclear segmentation. Cells were therefore segmented using trainable Weka segmentation [103] based on the diffuse Plo1-mCherry signal. Cytoplasmic concentrations were measured by manually placing two ROIs into the cytoplasm, one on each side of the nucleus. Plo1 enrichment at the SPB region and Cdc13 nuclear intensity were estimated by calculating the mean of the 40 brightest pixels in the cell. For the experiment in Figure 7 and S7, cells were segmented manually. Plo1 enrichment at the SPB region and nuclear Cdc13-sfGFP were estimated by calculating the mean of the 40 brightest pixels in the cell. Raw data curves were smoothed using the loess function in R (span 0.3).

The Pomegranate image analysis pipeline [104] was used for the 3D quantification of Cdc13-sfGFP shown in Fig. S1.

## Acknowledgments

We are grateful to Liv Erickson for experimental help, Yoshinori Watanabe for strains, and Doug Weidemann for comments.

## Author contributions

Conceptualization, SGC, JMR, SH; Investigation and Formal Analysis, SGC (Fig. 2A, 5, 6, S2, S6), JMR (Fig. 1, S1), WW (Fig. 2B), DV (Fig. 3, S3, S4), SH (Fig. 4, 7, S7); Software, VG, SJR; Writing – Original Draft, SH; Writing – Review & Editing, SGC, WW; Supervision, SH, SJR; Funding Acquisition, SH, SJR.

## Funding

Research reported in this publication was supported by the National Institute of General Medical Sciences of the National Institutes of Health under award numbers R35GM119723 and R35GM149565 (S.H.); V.G. was supported by SNSF grants CRSK-3_190526, 310030_204938, and CRSK-3_221036 awarded to S.J.R.

## Supplementary Material for

**Figure S1.**
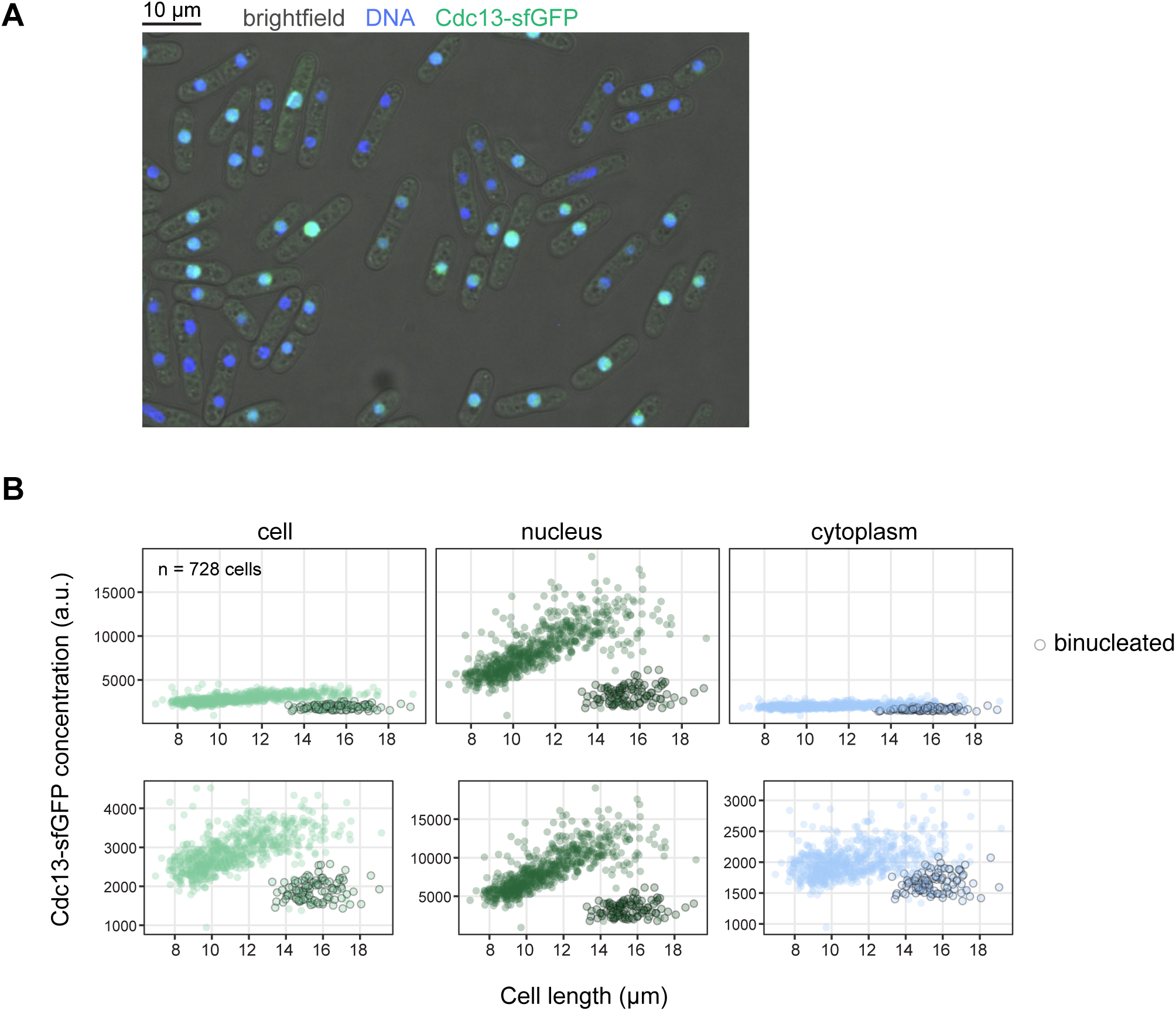
Analysis of Cdc13-sfGFP concentrations from a single time point of an asynchronous culture. **(A)** Example image of Cdc13-sfGFP-expressing cells, showing the central slice of a z-stack. **(B)** Cdc13-sfGFP concentrations relative to cell length, analyzed from images such as in (A). Cells and nuclei were segmented in 3D. Concentrations were calculated as the integrated signal intensities divided by volume for each compartment (cell, nucleus, cytoplasm). Each dot represents a single cell (n = 728). Graphs on top use the same y-axis scale; graphs at the bottom use a separate scale for each compartment. Binucleated cells are indicated by gray circles. In *S. pombe*, cells are generally binucleated in late mitosis, G1, and S phase.

**Figure S2.**
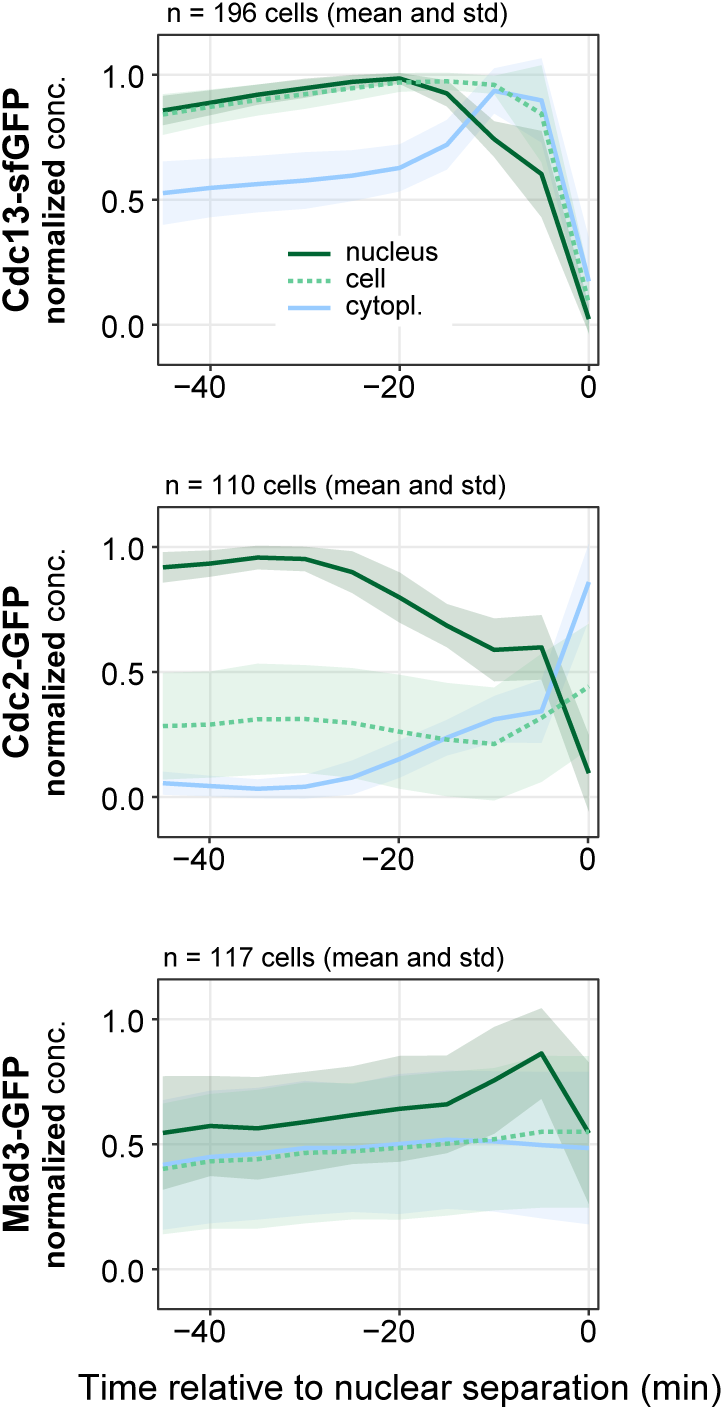
Normalized GFP concentrations for the experiment in. Figure 2. Live-cell imaging of Cdc13-sfGFP, Cdc2-GFP, and Mad3-GFP, all expressed from their endogenous genetic locus. Images were taken every 5 min. Same experiment as in Fig. 2 but normalized GFP concentrations are shown. GFP concentrations in cell, nucleus, and cytoplasm were normalized to the minimum and maximum signal within one cell cycle for each cell. Mean (lines) and standard deviation (area) across all quantified cells are shown.

**Figure S3.**
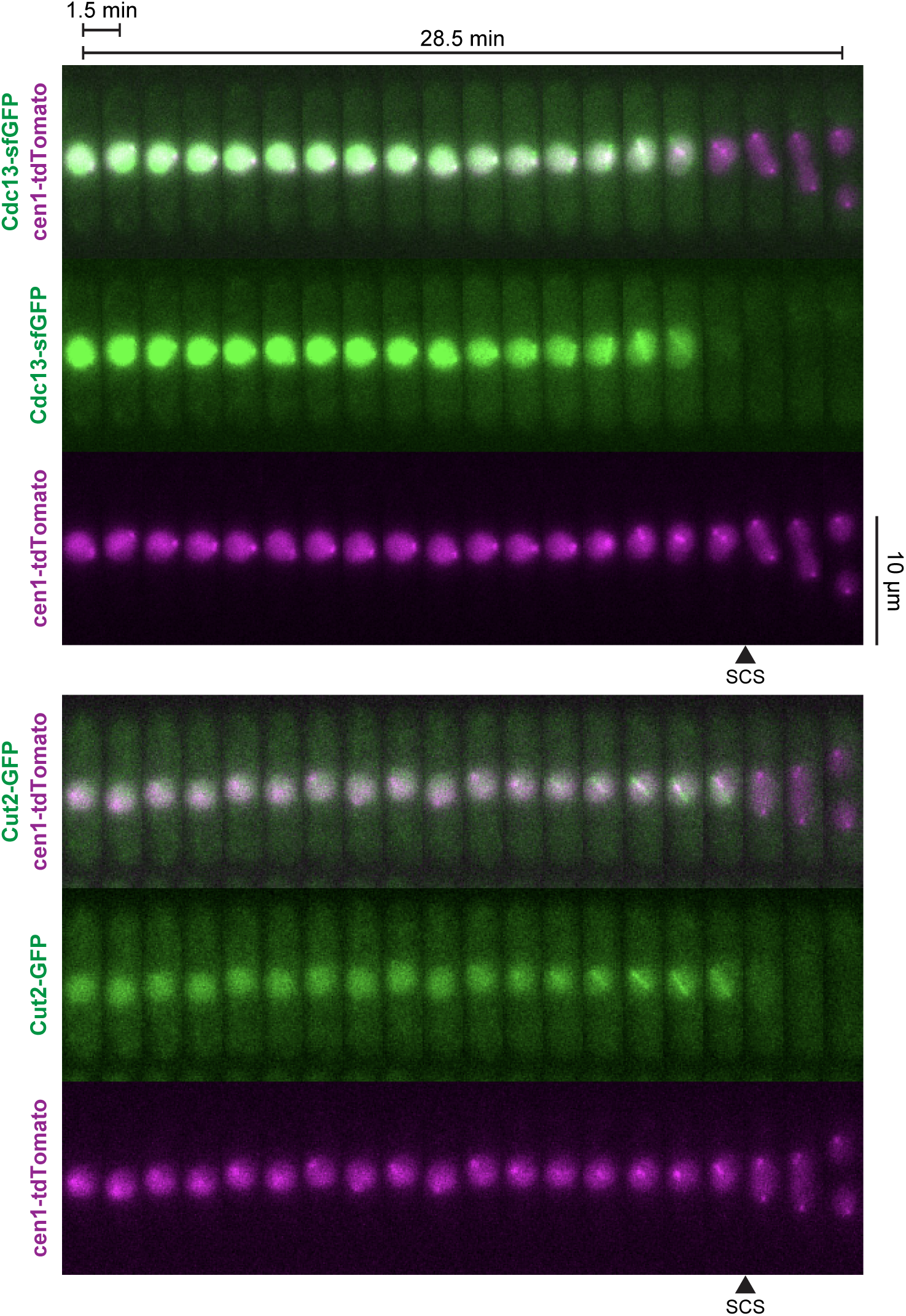
Cdc13 export and Cut2 import relative to sister chromatid separation. Live-cell imaging of Cdc13-sfGFP and Cut2-GFP, both expressed from their endogenous genetic locus. Images were taken every 15 sec; the kymographs only show every 6th image. Same cells as in Fig. 3, but additionally showing cen1-tdTomato (cen1-tetO/TetR-tdTomato), which was used to segment the nucleus and determine the time point of sister chromatid separation (SCS).

**Figure S4.**
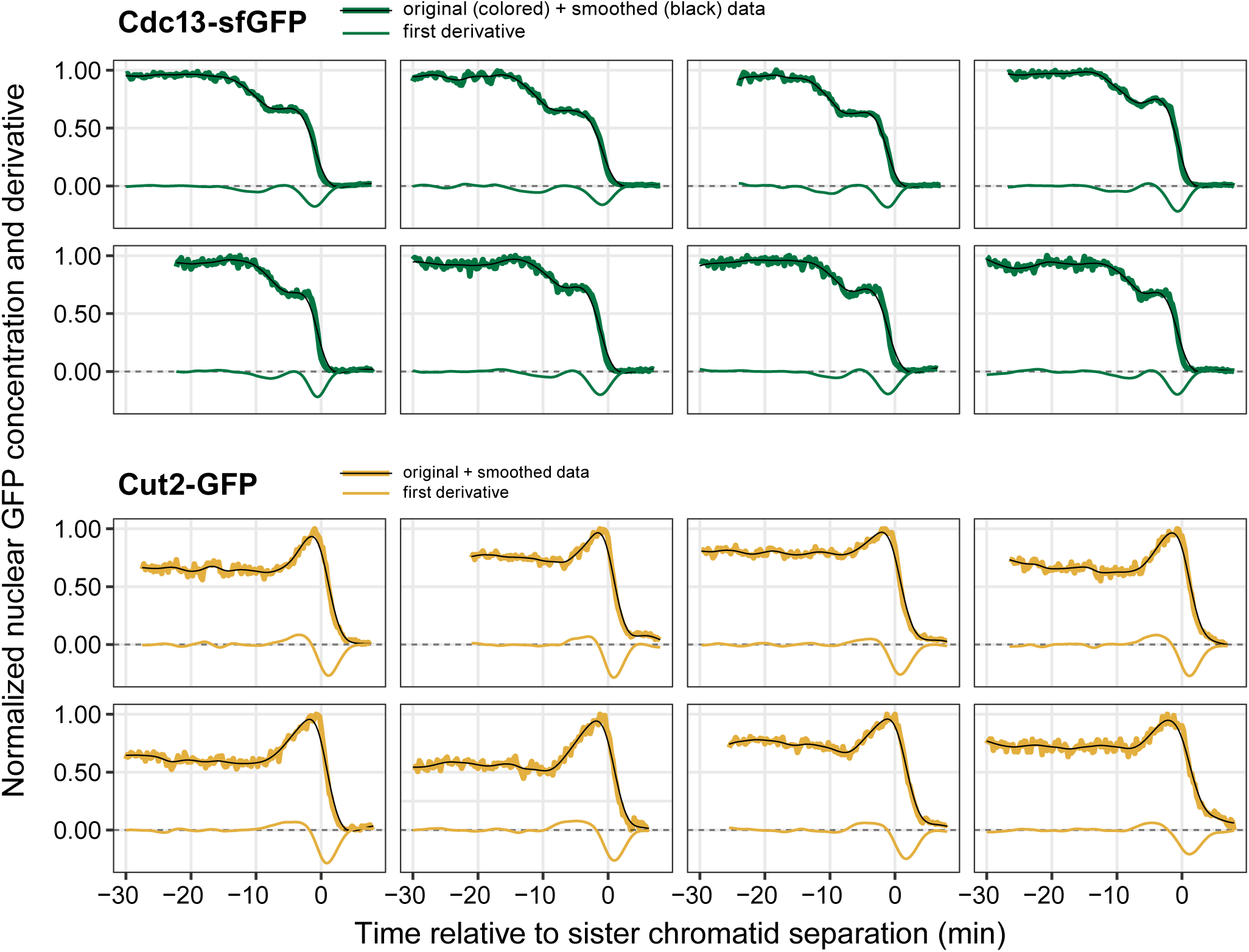
Cdc13 export and Cut2 import in individual cells. Live-cell imaging of Cdc13-sfGFP and Cut2-GFP, both expressed from their endogenous genetic locus. Images were taken every 15 sec. Same experiment as in Fig. 3, but showing data for individual cells to demonstrate the abrupt stop of Cdc13 export from the nucleus and onset of Cut2 import into the nucleus. Nuclear GFP concentrations were normalized to the maximum and minimum for each cell. Data smoothed using cubic splines (spar = 0.5). Black lines: smoothed data; colored thick lines: original data; colored thin lines: first derivative.

**Figure S5.**
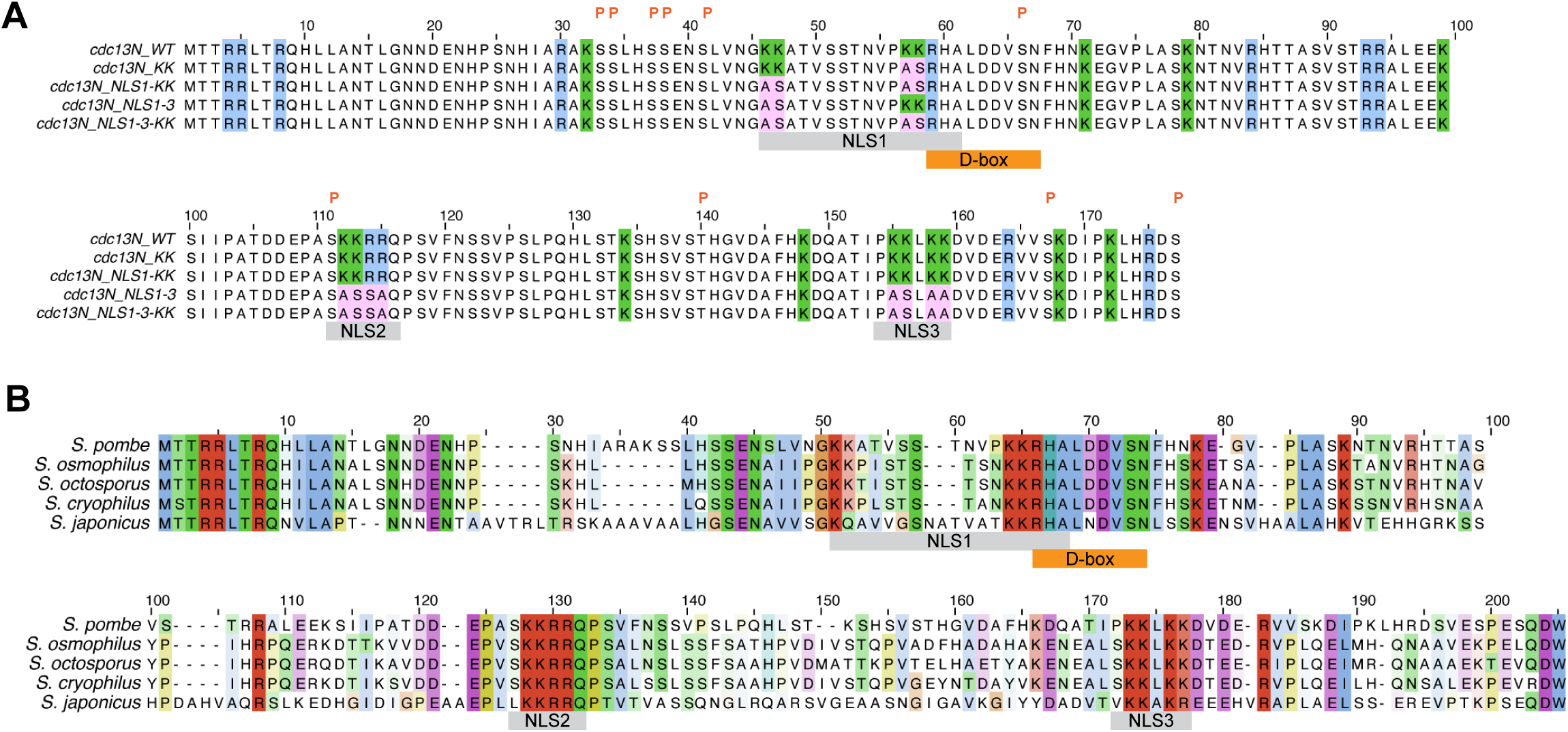
Amino acid sequence of the Cdc13 N-terminus. **(A)** Amino acid sequence of the *S. pombe* Cdc13 N-terminus with highlighting of arginine residues (blue), lysine residues (green), predicted NLSs (gray) and the D-box (orange). Residues mutated in different constructs are highlighted in pink. An orange “P” above the sequence indicates residues that have been found phosphorylated in mass spectrometry studies. **(B)** Alignment of the Cdc13 N-terminus from *S. pombe*, *S. osmophilus*, *S. octosporus*, *S. cryophilus*, and *S. japonicus*, showing conservation of the predicted NLSs.

**Figure S6.**
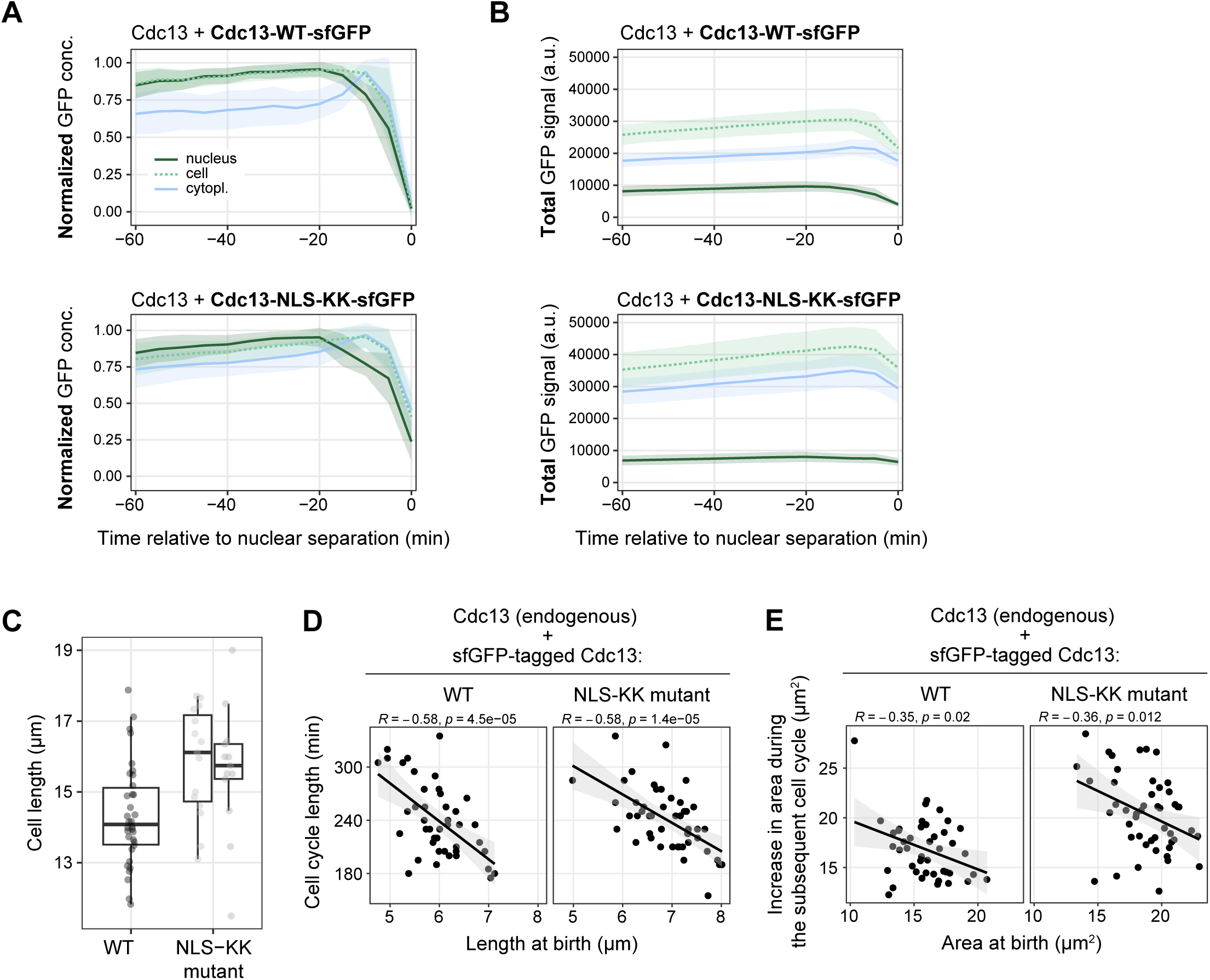
A fraction of Cdc13-NLS-KK is still exported from the nucleus prior to its degradation. Additional analysis from the live-cell imaging experiment in Fig. 6. Fluorescently (sfGFP) tagged wild-type (WT) or NLS-KK mutant Cdc13 were expressed from the exogenous *leu1* locus under the *cdc13* promoter. Untagged *cdc13* is expressed from its endogenous locus. **(A)** GFP concentrations were normalized to the minimum and maximum signal within one cell cycle to better demonstrate the pattern in whole cell and cytoplasmic GFP concentration. Mean (line) and standard deviation (area) from 60 and 57 cells, respectively. Curves are aligned to nuclear separation in anaphase (time 0). **(B)** The integrated, total signal in different regions (mean and standard deviation) is shown rather than the concentration. **(C)** Cell length of binucleated cells (i.e., just prior to cell division) from the experiment in Fig. 5. Two independent strains for the NSL-KK mutant; n = 35, 15, 13 cells. Individual cells (circles) and summary data; box: median and quartiles; whiskers: 1.5-times interquartile range. **(D)** Cell length at birth versus length of the subsequent cell cycle from the live-cell imaging experiment in Fig. 6. Same as Fig. 6D, except that one wild-type outlier cell with a long cell cycle length was removed. **(E)** Segmented cell area at birth versus the area increase during the subsequent cell cycle measured in the live-cell imaging experiment (Fig. 6).

**Figure S7.**
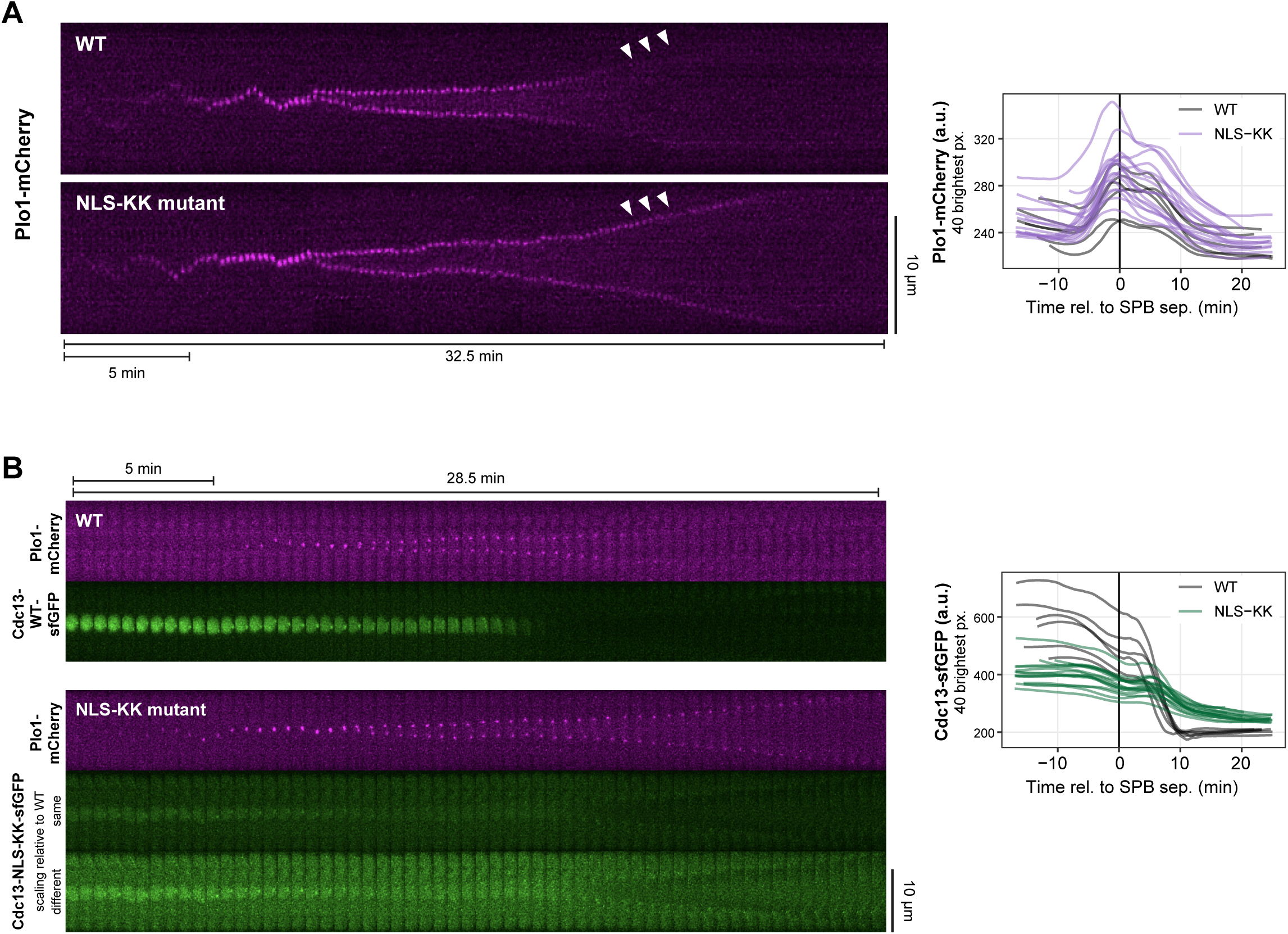
Plo1 removal from SPBs and Cdc13 degradation in cells expressing Cdc13-NLS-KK. Live-cell imaging of Plo1-mCherry, expressed from its endogenous locus, and sfGFP-tagged wild-type Cdc13 or NLS-KK mutant, expressed from the exogenous *leu1* locus under the *cdc13* promoter. Same experiment as in Fig. 7 (n = 6 and n = 16 cells from two independent strains, respectively). Images were taken every 15 sec. **(A)** Left: Kymographs centered on Plo1 localizing at SPBs in cells expressing wild-type Cdc13 or the NLS-KK mutant. We consistently observed that Plo1 retained the localization for longer (arrowheads) in the NLS-KK mutant cells. Right: Quantification of Plo1-mCherry in these cells, using the 40 brightest pixels in the cell as a proxy for Plo1 enrichment at SPBs. **(B)** Left: Kymographs from exemplary cells showing every 2nd image (every 30 sec). Cdc13-NLS-KK-sfGFP is shown once with the same scaling as Cdc13-WT-sfGFP and once with a different scaling in order to better visualize the relation between nuclear and cytoplasmic signal in the NLS-KK mutant strain. Right: Quantification of Cdc13-sfGFP (WT and NLS-KK) in these cells, using the 40 brightest pixels in the cell either as a proxy for nuclear concentration (prior to and early in mitosis) or for signal remaining late in mitosis. Cdc13-NLS-KK-sfGFP is inefficiently and not completely degraded at exit from mitosis. This could be due to mutating lysine residues required for ubiquitination or could be a consequence of the cytoplasmic enrichment of Cdc13-NLS-KK, where it may be less efficiently targeted by the APC/C (the E3 ligase ubiquitinating Cdc13), which is enriched in the nucleus.

**Supplementary Movie 1**

Live-cell imaging of an overexpressed N-terminal, sfGFP-tagged Cdc13 fragment (Cdc13-(1-177)-sfGFP; strain SU425’).

**Supplementary Movie 2**

Same cells as Supplementary Movie 1, but false-colored.

**Supplementary Movie 3**

Live-cell imaging of Cdc13-sfGFP (strain ST961).

**Supplementary Movie 4**

Same cell as Supplementary Movie 3, but false-colored.

**Supplementary Movie 5**

Same as Supplementary Movie 4, but the gamma value was adjusted to 0.7 to better visualize enrichment at SPBs and spindle during mitosis.

## Notes

### Competing Interest Statement

The authors have declared no competing interest.

### Summary of Updates

author affiliations updated; Figure 7 changed; existing Figure 7 moved to Figure 8; text changes to reflect added figure; Supplemental Movies added

## References

1. Hochegger H, Takeda S, Hunt T. 2008 Cyclin-dependent kinases and cell-cycle transitions: does one fit all? Nat. Rev. Mol. Cell Biol. 9, 910–916. (doi:10.1038/nrm2510)

2. Malumbres M. 2014 Cyclin-dependent kinases. Genome Biol 15, 122. (doi:10.1186/gb4184)

3. Örd M, Loog M. 2019 How the cell cycle clock ticks. Mol. Biol. Cell 30, 169–172. (doi:10.1091/mbc.e18-05-0272)

4. Moser BA, Russell P. 2000 Cell cycle regulation in Schizosaccharomyces pombe. Curr Opin Microbiol 3, 631–636. (doi:10.1016/s1369-5274(00)00152-1)

5. Fisher DL, Nurse P. 1996 A single fission yeast mitotic cyclin B p34cdc2 kinase promotes both S-phase and mitosis in the absence of G1 cyclins. EMBO J 15, 850–860. (doi:10.1002/j.1460-2075.1996.tb00420.x)

6. Coudreuse D, Nurse P. 2010 Driving the cell cycle with a minimal CDK control network. Nature 468, 1074–1079. (doi:10.1038/nature09543)

7. Alfa CE, Booher R, Beach D, Hyams JS. 1989 Fission yeast cyclin: subcellular localisation and cell cycle regulation. J. Cell Sci. 1989, 9–19. (doi:10.1242/jcs.1989.supplement_12.2)

8. Alfa CE, Booher RN, Beach DH, Hyams JS. 1989 The fission yeast cdc2/cdc13/suc1 protein kinase: Regulation of catalytic activity and nuclear localization. Cell 58, 485–497. (doi:10.1016/0092-8674(89)90429-7)

9. Moreno S, Hayles J, Nurse P. 1989 Regulation of p34cdc2 protein kinase during mitosis. Cell 58, 361–372. (doi:10.1016/0092-8674(89)90850-7)

10. Yamano H, Gannon J, Hunt T. 1996 The role of proteolysis in cell cycle progression in Schizosaccharomyces pombe. EMBO J. 15, 5268–5279. (doi:10.1002/j.1460-2075.1996.tb00912.x)

11. Yamano H, Tsurumi C, Gannon J, Hunt T. 1998 The role of the destruction box and its neighbouring lysine residues in cyclin B for anaphase ubiquitin-dependent proteolysis in fission yeast: defining the D-box receptor. EMBO J. 17, 5670–5678. (doi:10.1093/emboj/17.19.5670)

12. Decottignies A, Zarzov P, Nurse P. 2001 In vivo localisation of fission yeast cyclin-dependent kinase cdc2p and cyclin B cdc13p during mitosis and meiosis. J. Cell Sci. 114, 2627–2640. (doi:10.1242/jcs.114.14.2627)

13. Pines J. 1999 Four-dimensional control of the cell cycle. Nat. Cell Biol. 1, E73–E79. (doi:10.1038/11041)

14. Alfa CE, Ducommun B, Beach D, Hyams JS. 1990 Distinct nuclear and spindle pole body populations of cyclin–cdc2 in fission yeast. Nature 347, 680–682. (doi:10.1038/347680a0)

15. Yanagida M, Yamashita YM, Tatebe H, Ishii K, Kumada K, Nakaseko Y. 1999 Control of metaphaseanaphase progression by proteolysis: cyclosome function regulated by the protein kinase A pathway, ubiquitination and localization. Philos. Trans. R. Soc. Lond. Ser. B: Biol. Sci. 354, 1559–1570. (doi:10.1098/rstb.1999.0499)

16. Tatebe H, Goshima G, Takeda K, Nakagawa T, Kinoshita K, Yanagida M. 2001 Fission yeast living mitosis visualized by GFP-tagged gene products. Micron 32, 67–74. (doi:10.1016/s0968-4328(00)00023-8)

17. Basu S, Roberts EL, Jones AW, Swaffer MP, Snijders AP, Nurse P. 2020 The Hydrophobic Patch Directs Cyclin B to Centrosomes to Promote Global CDK Phosphorylation at Mitosis. Curr. Biol. 30, 883–892.e4. (doi:10.1016/j.cub.2019.12.053)

18. Curran S, Dey G, Rees P, Nurse P. 2022 A quantitative and spatial analysis of cell cycle regulators during the fission yeast cycle. Proc. Natl. Acad. Sci. 119, e2206172119. (doi:10.1073/pnas.2206172119)

19. Roberts EL, Greenwood J, Kapadia N, Auchynnikava T, Basu S, Nurse P. 2024 CDK activity at the centrosome regulates the cell cycle. Cell Rep. 43, 114066. (doi:10.1016/j.celrep.2024.114066)

20. Alfa CE, Gallagher IM, Hyams JS. 1991 Subcellular Localization of the p34cdc2/p63cdc13 Protein Kinase in Fission Yeast. Cold Spring Harb. Symp. Quant. Biol. 56, 489–494. (doi:10.1101/sqb.1991.056.01.055)

21. Gallagher IM, Alfa CE, Hyams JS. 1993 p63cdc13, a B-type cyclin, is associated with both the nucleolar and chromatin domains of the fission yeast nucleus. Mol. Biol. Cell 4, 1087–1096. (doi:10.1091/mbc.4.11.1087)

22. Ren L, Feoktistova A, McDonald WH, Haese GD, Morrell JL, Gould KL. 2005 Analysis of the Role of Phosphorylation in Fission Yeast Cdc13p/CyclinB Function. J Biol Chem 280, 14591–14596. (doi:10.1074/jbc.m500560200)

23. Hiraoka M, Kiyota Y, Kawai S, Notsu Y, Yamada K, Kurashima K, Chang J-W, Shimazaki S, Yamamoto A. 2023 CDK actively contributes to establishment of the stationary phase state in fission yeast. J. Cell Sci. 136. (doi:10.1242/jcs.260727)

24. Pines J, Hunter T. 1991 Human cyclins A and B1 are differentially located in the cell and undergo cell cycle-dependent nuclear transport. J. cell Biol. 115, 1–17. (doi:10.1083/jcb.115.1.1)

25. Bailly E, Pines J, Hunter T, Bornens M. 1992 Cytoplasmic accumulation of cyclin B1 in human cells: Association with a detergent-resistant compartment and with the centrosome. J. Cell Sci. 101, 529–545. (doi:10.1242/jcs.101.3.529)

26. Elzen N den, Pines J. 2001 Cyclin a Is Destroyed in Prometaphase and Can Delay Chromosome Alignment and Anaphase. J. Cell Biol. 153, 121–136. (doi:10.1083/jcb.153.1.121)

27. Jackman M, Kubota Y, Elzen N den, Hagting A, Pines J. 2002 Cyclin A-and Cyclin E-Cdk Complexes Shuttle between the Nucleus and the Cytoplasm. Mol. Biol. Cell 13, 1030–1045. (doi:10.1091/mbc.01-07-0361)

28. Pascreau G, Eckerdt F, Churchill MEA, Maller JL. 2010 Discovery of a distinct domain in cyclin A sufficient for centrosomal localization independently of Cdk binding. Proc. Natl. Acad. Sci. 107, 2932–2937. (doi:10.1073/pnas.0914874107)

29. Boer LD, Oakes V, Beamish H, Giles N, Stevens F, Somodevilla-Torres M, DeSouza C, Gabrielli B. 2008 Cyclin A/cdk2 coordinates centrosomal and nuclear mitotic events. Oncogene 27, 4261–4268. (doi:10.1038/onc.2008.74)

30. Zerjatke T, Gak IA, Kirova D, Fuhrmann M, Daniel K, Gonciarz M, Müller D, Glauche I, Mansfeld J. 2017 Quantitative Cell Cycle Analysis Based on an Endogenous All-in-One Reporter for Cell Tracking and Classification. Cell Rep. 19, 1953–1966. (doi:10.1016/j.celrep.2017.05.022)

31. Cascales HS, Burdova K, Middleton A, Kuzin V, Müllers E, Stoy H, Baranello L, Macurek L, Lindqvist A. 2021 Cyclin A2 localises in the cytoplasm at the S/G2 transition to activate PLK1. Life Sci. Alliance 4, e202000980. (doi:10.26508/lsa.202000980)

32. Clute P, Pines J. 1999 Temporal and spatial control of cyclin B1 destruction in metaphase. Nat. Cell Biol. 1, 82–87. (doi:10.1038/10049)

33. Hagting A, Jackman M, Simpson K, Pines J. 1999 Translocation of cyclin B1 to the nucleus at prophase requires a phosphorylation-dependent nuclear import signal. Curr. Biol. 9, 680–689. (doi:10.1016/s0960-9822(99)80308-x)

34. Gavet O, Pines J. 2010 Activation of cyclin B1–Cdk1 synchronizes events in the nucleus and the cytoplasm at mitosis. J. Cell Biol. 189, 247–259. (doi:10.1083/jcb.200909144)

35. Gavet O, Pines J. 2010 Progressive Activation of CyclinB1-Cdk1 Coordinates Entry to Mitosis. Dev. Cell 18, 533–543. (doi:10.1016/j.devcel.2010.02.013)

36. Santos SDM, Wollman R, Meyer T, Ferrell JE. 2012 Spatial positive feedback at the onset of mitosis. Cell 149, 1500–13. (doi:10.1016/j.cell.2012.05.028)

37. Ferrell JE. 1998 How regulated protein translocation can produce switch-like responses. Trends Biochem. Sci. 23, 461–465. (doi:10.1016/s0968-0004(98)01316-4)

38. Bentley AM, Normand G, Hoyt J, King RW. 2007 Distinct Sequence Elements of Cyclin B1 Promote Localization to Chromatin, Centrosomes, and Kinetochores during Mitosis. Mol. Biol. Cell 18, 4847–4858. (doi:10.1091/mbc.e06-06-0539)

39. Chen Q, Zhang X, Jiang Q, Clarke PR, Zhang C. 2008 Cyclin B1 is localized to unattached kinetochores and contributes to efficient microtubule attachment and proper chromosome alignment during mitosis. Cell Res. 18, 268–280. (doi:10.1038/cr.2008.11)

40. Alfonso-Pérez T, Hayward D, Holder J, Gruneberg U, Barr FA. 2019 MAD1-dependent recruitment of CDK1-CCNB1 to kinetochores promotes spindle checkpoint signaling. J. Cell Biol., jcb.201808015. (doi:10.1083/jcb.201808015)

41. Allan LA, Reis MC, Ciossani G, Veld PJH in‘t, Wohlgemuth S, Kops GJ, Musacchio A, Saurin AT. 2020 Cyclin B1 scaffolds MAD1 at the kinetochore corona to activate the mitotic checkpoint. EMBO J. 39, EMBJ2019103180. (doi:10.15252/embj.2019103180)

42. Kapadia N, Nurse P. 2025 Spatiotemporal orchestration of mitosis by cyclin-dependent kinase. Nature 643, 1391–1399. (doi:10.1038/s41586-025-09172-y)

43. Tallada VA, Tanaka K, Yanagida M, Hagan IM. 2009 The S. pombe mitotic regulator Cut12 promotes spindle pole body activation and integration into the nuclear envelope. J. Cell Biol. 185, 875–888. (doi:10.1083/jcb.200812108)

44. Fong CS, Sato M, Toda T. 2010 Fission yeast Pcp1 links polo kinase-mediated mitotic entry to γ-tubulin-dependent spindle formation. EMBO J. 29, 120–130. (doi:10.1038/emboj.2009.331)

45. Masuda H, Fong CS, Ohtsuki C, Haraguchi T, Hiraoka Y. 2011 Spatiotemporal regulations of Wee1 at the G2/M transition. Mol. Biol. Cell 22, 555–569. (doi:10.1091/mbc.e10-07-0644)

46. Fernández-Álvarez A, Bez C, O’Toole ET, Morphew M, Cooper JP. 2016 Mitotic Nuclear Envelope Breakdown and Spindle Nucleation Are Controlled by Interphase Contacts between Centromeres and the Nuclear Envelope. Dev. Cell 39, 544–559. (doi:10.1016/j.devcel.2016.10.021)

47. Bestul AJ, Yu Z, Unruh JR, Jaspersen SL. 2021 Redistribution of centrosomal proteins by centromeres and Polo kinase controls partial nuclear envelope breakdown in fission yeast. Mol. Biol. Cell 32, 1487–1500. (doi:10.1091/mbc.e21-05-0239)

48. Sugiyama H, Goto Y, Kondo Y, Coudreuse D, Aoki K. 2024 Live-cell imaging defines a threshold in CDK activity at the G2/M transition. Dev. Cell 59, 545–557.e4. (doi:10.1016/j.devcel.2023.12.014)

49. Smoyer CJ, Jaspersen SL. 2014 Breaking down the wall: the nuclear envelope during mitosis. Curr. Opin. Cell Biol. 26, 1–9. (doi:10.1016/j.ceb.2013.08.002)

50. Rüthnick D, Schiebel E. 2018 Duplication and Nuclear Envelope Insertion of the Yeast Microtubule Organizing Centre, the Spindle Pole Body. Cells 7, 42. (doi:10.3390/cells7050042)

51. Kobayashi H, Stewart E, Poon R, Adamczewski JP, Gannon J, Hunt T. 1992 Identification of the domains in cyclin A required for binding to, and activation of, p34cdc2 and p32cdk2 protein kinase subunits. Mol. Biol. Cell 3, 1279–1294. (doi:10.1091/mbc.3.11.1279)

52. Lees EM, Harlow E. 1993 Sequences Within the Conserved Cyclin Box of Human Cyclin A are Sufficient for Binding to and Activation of cdc2 Kinase. Mol. Cell. Biol. 13, 1194–1201. (doi:10.1128/mcb.13.2.1194-1201.1993)

53. Hagan IM. 2008 The spindle pole body plays a key role in controlling mitotic commitment in the fission yeast Schizosaccharomyces pombe. Biochem. Soc. Trans. 36, 1097–1101. (doi:10.1042/bst0361097)

54. Archambault V, Glover DM. 2009 Polo-like kinases: conservation and divergence in their functions and regulation. Nat. Rev. Mol. Cell Biol. 10, 265–275. (doi:10.1038/nrm2653)

55. Lindqvist A, Rodríguez-Bravo V, Medema RH. 2009 The decision to enter mitosis: feedback and redundancy in the mitotic entry network. J. Cell Biol. 185, 193–202. (doi:10.1083/jcb.200812045)

56. Pines J, Hagan I. 2011 The Renaissance or the cuckoo clock. Philos. Trans. R. Soc. B: Biol. Sci. 366, 3625–3634. (doi:10.1098/rstb.2011.0080)

57. Hagan IM, Grallert A. 2013 Spatial control of mitotic commitment in fission yeast. Biochem. Soc. Trans. 41, 1766–1771. (doi:10.1042/bst20130190)

58. Bähler J, Steever AB, Wheatley S, Wang Y, Pringle JR, Gould KL, McCollum D. 1998 Role of Polo Kinase and Mid1p in Determining the Site of Cell Division in Fission Yeast. J. Cell Biol. 143, 1603–1616. (doi:10.1083/jcb.143.6.1603)

59. Mulvihill DP, Petersen J, Ohkura H, Glover DM, Hagan IM. 1999 Plo1 Kinase Recruitment to the Spindle Pole Body and Its Role in Cell Division in Schizosaccharomyces pombe. Mol. Biol. Cell 10, 2771–2785. (doi:10.1091/mbc.10.8.2771)

60. MacIver FH, Tanaka K, Robertson AM, Hagan IM. 2003 Physical and functional interactions between polo kinase and the spindle pole component Cut12 regulate mitotic commitment in S. pombe. Genes Dev. 17, 1507–1523. (doi:10.1101/gad.256003)

61. Grallert A, Patel A, Tallada VA, Chan KY, Bagley S, Krapp A, Simanis V, Hagan IM. 2013 Centrosomal MPF triggers the mitotic and morphogenetic switches of fission yeast. Nat. Cell Biol. 15, 88–95. (doi:10.1038/ncb2633)

62. Ding R, West RR, Morphew DM, Oakley BR, McIntosh JR. 1997 The spindle pole body of Schizosaccharomyces pombe enters and leaves the nuclear envelope as the cell cycle proceeds. Mol. Biol. Cell 8, 1461–1479. (doi:10.1091/mbc.8.8.1461)

63. West RR, Vaisberg EV, Ding R, Nurse P, McIntosh JR. 1998 cut11 +: A Gene Required for Cell Cycle-dependent Spindle Pole Body Anchoring in the Nuclear Envelope and Bipolar Spindle Formation in Schizosaccharomyces pombe. Mol. Biol. Cell 9, 2839–2855. (doi:10.1091/mbc.9.10.2839)

64. Yang J, Bardes ESG, Moore JD, Brennan J, Powers MA, Kornbluth S. 1998 Control of Cyclin B1 localization through regulated binding of the nuclear export factor CRM1. Genes Dev. 12, 2131–2143. (doi:10.1101/gad.12.14.2131)

65. Hagting A, Karlsson C, Clute P, Jackman M, Pines J. 1998 MPF localization is controlled by nuclear export. EMBO J. 17, 4127–4138. (doi:10.1093/emboj/17.14.4127)

66. Toyoshima F, Moriguchi T, Wada A, Fukuda M, Nishida E. 1998 Nuclear export of cyclin B1 and its possible role in the DNA damage-induced G2 checkpoint. EMBO J 17, 2728–2735. (doi:10.1093/emboj/17.10.2728)

67. Jin P, Hardy S, Morgan DO. 1998 Nuclear Localization of Cyclin B1 Controls Mitotic Entry After DNA Damage. J. Cell Biol. 141, 875–885. (doi:10.1083/jcb.141.4.875)

68. Takizawa CG, Weis K, Morgan DO. 1999 Ran-independent nuclear import of cyclin B1– Cdc2 by importin β. Proc. Natl. Acad. Sci. 96, 7938–7943. (doi:10.1073/pnas.96.14.7938)

69. Bendris N, Lemmers B, Blanchard J-M, Arsic N. 2011 Cyclin A2 Mutagenesis Analysis: A New Insight into CDK Activation and Cellular Localization Requirements. PLoS ONE 6, e22879. (doi:10.1371/journal.pone.0022879)

70. Moore JD, Yang J, Truant R, Kornbluth S. 1999 Nuclear Import of Cdk/Cyclin Complexes: Identification of Distinct Mechanisms for Import of Cdk2/Cyclin E and Cdc2/Cyclin B1. J. Cell Biol. 144, 213–224. (doi:10.1083/jcb.144.2.213)

71. Rhind N. 2021 Cell-size control. Curr. Biol. 31, R1414–R1420. (doi:10.1016/j.cub.2021.09.017)

72. Lopez-Girona A, Kanoh J, Russell P. 2001 Nuclear exclusion of Cdc25 is not required for the DNA damage checkpoint in fission yeast. Curr. Biol. 11, 50–54. (doi:10.1016/s0960-9822(00)00026-9)

73. Chua G, Lingner C, Frazer C, Young PG. 2002 The sal3+ Gene Encodes an Importin-β Implicated in the Nuclear Import of Cdc25 in Schizosaccharomyces pombe. Genetics 162, 689– 703. (doi:10.1093/genetics/162.2.689)

74. Ferrell JE, Ha SH. 2014 Ultrasensitivity part I: Michaelian responses and zero-order ultrasensitivity. Trends Biochem. Sci. 39, 496–503. (doi:10.1016/j.tibs.2014.08.003)

75. Ferrell JE, Jr, Ha SH. 2014 Ultrasensitivity part II: multisite phosphorylation, stoichiometric inhibitors, and positive feedback. Trends Biochem. Sci. 39, 556–569. (doi:10.1016/j.tibs.2014.09.003)

76. MacCoss MJ et al. 2002 Shotgun identification of protein modifications from protein complexes and lens tissue. Proc. Natl. Acad. Sci. 99, 7900–7905. (doi:10.1073/pnas.122231399)

77. Unwin RD, Griffiths JR, Leverentz MK, Grallert A, Hagan IM, Whetton AD. 2005 Multiple Reaction Monitoring to Identify Sites of Protein Phosphorylation with High Sensitivity * S. Mol. Cell. Proteom. 4, 1134–1144. (doi:10.1074/mcp.m500113-mcp200)

78. Koch A, Krug K, Pengelley S, Macek B, Hauf S. 2011 Mitotic Substrates of the Kinase Aurora with Roles in Chromatin Regulation Identified Through Quantitative Phosphoproteomics of Fission Yeast. Sci. Signal. 4, rs6. (doi:10.1126/scisignal.2001588)

79. Carpy A, Krug K, Graf S, Koch A, Popic S, Hauf S, Macek B. 2014 Absolute Proteome and Phosphoproteome Dynamics during the Cell Cycle of Schizosaccharomyces pombe (Fission Yeast)*. Mol. Cell. Proteom. 13, 1925–1936. (doi:10.1074/mcp.m113.035824)

80. Tay YD, Leda M, Spanos C, Rappsilber J, Goryachev AB, Sawin KE. 2019 Fission Yeast NDR/LATS Kinase Orb6 Regulates Exocytosis via Phosphorylation of the Exocyst Complex. Cell Rep. 26, 1654–1667.e7. (doi:10.1016/j.celrep.2019.01.027)

81. Halova L et al. 2021 A TOR (target of rapamycin) and nutritional phosphoproteome of fission yeast reveals novel targets in networks conserved in humans. Open Biol 11, 200405. (doi:10.1098/rsob.200405)

82. Li J, Meyer AN, Donoghue DJ. 1997 Nuclear localization of cyclin B1 mediates its biological activity and is regulated by phosphorylation. Proc. Natl. Acad. Sci. 94, 502–507. (doi:10.1073/pnas.94.2.502)

83. Yang J, Song H, Walsh S, Bardes ESG, Kornbluth S. 2001 Combinatorial Control of Cyclin B1 Nuclear Trafficking through Phosphorylation at Multiple Sites*. J Biol Chem 276, 3604– 3609. (doi:10.1074/jbc.m008151200)

84. Toyoshima-Morimoto F, Taniguchi E, Shinya N, Iwamatsu A, Nishida E. 2001 Polo-like kinase 1 phosphorylates cyclin B1 and targets it to the nucleus during prophase. Nature 410, 215–220. (doi:10.1038/35065617)

85. Swaffer MP, Jones AW, Flynn HR, Snijders AP, Nurse P. 2016 CDK Substrate Phosphorylation and Ordering the Cell Cycle. Cell 167, 1750–1761.e16. (doi:10.1016/j.cell.2016.11.034)

86. Swaffer MP, Jones AW, Flynn HR, Snijders AP, Nurse P. 2018 Quantitative Phosphoproteomics Reveals the Signaling Dynamics of Cell-Cycle Kinases in the Fission Yeast Schizosaccharomyces pombe. Cell Rep. 24, 503–514. (doi:10.1016/j.celrep.2018.06.036)

87. Moris N, Shrivastava J, Jeffery L, Li J-J, Hayles J, Nurse P. 2016 A genome–wide screen to identify genes controlling the rate of entry into mitosis in fission yeast. Cell Cycle 15, 00–00. (doi:10.1080/15384101.2016.1242535)

88. Jackman M, Lindon C, Nigg EA, Pines J. 2003 Active cyclin B1–Cdk1 first appears on centrosomes in prophase. Nat. Cell Biol. 5, 143–148. (doi:10.1038/ncb918)

89. Cavanaugh AM, Jaspersen SL. 2016 Big Lessons from Little Yeast: Budding and Fission Yeast Centrosome Structure, Duplication, and Function. Annu. Rev. Genet. 51, 1–23. (doi:10.1146/annurev-genet-120116-024733)

90. Bridge AJ, Morphew M, Bartlett R, Hagan IM. 1998 The fission yeast SPB component Cut12 links bipolar spindle formation to mitotic control. Genes Dev. 12, 927–942. (doi:10.1101/gad.12.7.927)

91. Tamm T, Grallert A, Grossman EPS, Alvarez-Tabares I, Stevens FE, Hagan IM. 2011 Brr6 drives the Schizosaccharomyces pombe spindle pole body nuclear envelope insertion/extrusion cycle. J. Cell Biol. 195, 467–484. (doi:10.1083/jcb.201106076)

92. Ohkura H, Hagan IM, Glover DM. 1995 The conserved Schizosaccharomyces pombe kinase plo1, required to form a bipolar spindle, the actin ring, and septum, can drive septum formation in G1 and G2 cells. Genes Dev. 9, 1059–1073. (doi:10.1101/gad.9.9.1059)

93. Wälde S, King MC. 2014 The KASH protein Kms2 coordinates mitotic remodeling of the spindle pole body. J. Cell Sci. 127, 3625–3640. (doi:10.1242/jcs.154997)

94. Dischinger S, Krapp A, Xie L, Paulson JR, Simanis V. 2008 Chemical genetic analysis of the regulatory role of Cdc2p in the S. pombe septation initiation network. J. Cell Sci. 121, 843–853. (doi:10.1242/jcs.021584)

95. Tanaka K, Petersen J, MacIver F, Mulvihill DP, Glover DM, Hagan IM. 2001 The role of Plo1 kinase in mitotic commitment and septation in Schizosaccharomyces pombe. EMBO J. 20, 1259–1270. (doi:10.1093/emboj/20.6.1259)

96. Kamenz J, Mihaljev T, Kubis A, Legewie S, Hauf S. 2015 Robust Ordering of Anaphase Events by Adaptive Thresholds and Competing Degradation Pathways. Mol. Cell 60, 446–459. (doi:10.1016/j.molcel.2015.09.022)

97. Sakuno T, Tada K, Watanabe Y. 2009 Kinetochore geometry defined by cohesion within the centromere. Nature 458, 852–858. (doi:10.1038/nature07876)

98. Heinrich S, Windecker H, Hustedt N, Hauf S. 2012 Mph1 kinetochore localization is crucial and upstream in the hierarchy of spindle assembly checkpoint protein recruitment to kinetochores. J. Cell Sci. 125, 4720–4727. (doi:10.1242/jcs.110387)

99. Esposito E, Weidemann DE, Rogers JM, Morton CM, Baybay EK, Chen J, Hauf S. 2022 Mitotic checkpoint gene expression is tuned by codon usage bias. EMBO J. 41, EMBJ2021107896. (doi:10.15252/embj.2021107896)

100. Matsuyama A, Shirai A, Yashiroda Y, Kamata A, Horinouchi S, Yoshida M. 2004 pDUAL, a multipurpose, multicopy vector capable of chromosomal integration in fission yeast. Yeast 21, 1289–1305. (doi:10.1002/yea.1181)

101. Dietler N et al. 2020 A convolutional neural network segments yeast microscopy images with high accuracy. Nat. Commun. 11, 5723. (doi:10.1038/s41467-020-19557-4)

102. Schindelin J et al. 2012 Fiji: an open-source platform for biological-image analysis. Nat. Methods 9, 676–682. (doi:10.1038/nmeth.2019)

103. Arganda-Carreras I, Kaynig V, Rueden C, Eliceiri KW, Schindelin J, Cardona A, Seung HS. 2017 Trainable Weka Segmentation: a machine learning tool for microscopy pixel classification. Bioinformatics 33, 2424–2426. (doi:10.1093/bioinformatics/btx180)

104. Baybay EK, Esposito E, Hauf S. 2020 Pomegranate: 2D segmentation and 3D reconstruction for fission yeast and other radially symmetric cells. Sci. Rep. 10, 16580. (doi:10.1038/s41598-020-73597-w)

