## Supplementary figures and images for "A distinct phase of cyclin B (Cdc13) nuclear export at mitotic entry in *S. pombe*"

### Supplemental Movie 1

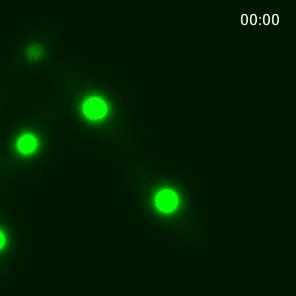

### Supplemental Movie 2

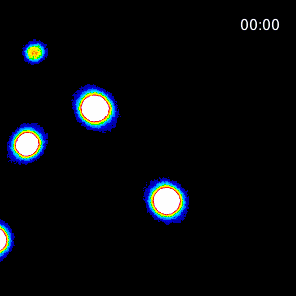

### Supplemental Movie 3

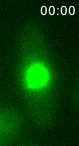

### Supplemental Movie 4

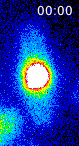

### Supplemental Movie 5

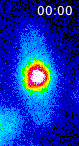
